# Normalizing single-cell RNA sequencing data with internal spike-in-like genes

**DOI:** 10.1101/2020.07.10.198077

**Authors:** Li Lin, Minfang Song, Yong Jiang, Xiaojing Zhao, Haopeng Wang, Liye Zhang

## Abstract

Normalization with respect to sequencing depth is a crucial step in single-cell RNA sequencing preprocessing. Most methods normalize data using the whole transcriptome based on the assumption that the majority of transcriptome remains constant and are unable to detect drastic changes of the transcriptome. Here, we develop an algorithm based on a small fraction of constantly expressed genes as internal spike-ins to normalize single cell RNA sequencing data. We demonstrate that the transcriptome of single cells may undergo drastic changes in several case study datasets and accounting for such heterogeneity by ISnorm improves the performance of downstream analyzes.

## INTRODUCTION

Single-cell RNA sequencing (scRNA-seq) provides researchers with a powerful tool to investigate questions that cannot be addressed by bulk sequencing. The scRNA-seq data shares similar features with data from bulk RNA-Seq such as over dispersion of gene expression, but also has several distinct features, such as high sparsity (i.e. high proportion of zero read counts in the data) (1). These features can be derived from both technical noises and biological variations, which provide challenges for computational methods to handle scRNA-seq data. Among the computational methods, normalization is one of the most important steps in scRNA-seq data preprocessing and exerts significant effect on downstream analyses. As current high-throughput sequencing techniques provides compositional data, where the value of one feature is a proportion and is only meaningful when compared to other features, normalization serves the function of transforming relative abundances into absolute abundances and making the data interpretable by conventional statistical methods (2-4). Although there are many existing methods of normalization, most of them adopt the principle of normalization to effective library size, which is very similar to the centered log-ratio transformation (clr) strategy (3). They calculate a cell-specific scaling factor (size factor) and divide raw counts from each cell by its size factor to account for the difference of RNA capture efficiency, sequencing depth or other potential technical biases between individual cells. Several state-of-the-art methods have been developed to better handle specific technical biases in single cell RNA-Seq technology such as dropout effects (2-4). However, almost all methods assume that the majority of the transcriptome remains constant and seek to minimize the number of differentially expressed (DE) genes. Therefore, systematic biases may be introduced when the transcriptome undergoes drastic changes.

In RNA sequencing, to capture the drastic changes in transcriptome, a set of synthetic control transcripts (external spike-ins) are normally used for normalization (5). The same amounts of external spike-in RNAs are added to each sample (bulk samples or single cells) serve as external references. The usage of external spike-ins is based on the assumption that all technical factors affect extrinsic and intrinsic genes in the same manner (6). However, there are several limitations on the adoption of external spike-ins in scRNA-seq (7) (e.g. too many spike-ins overwhelm signals from intrinsic genes; external spike-ins are not always available; differences in cell lysis efficiency). Most importantly, external spike-ins can vary significantly even between technical replicates (6).

Considering the potential caveats of external spike-ins, normalization with an internal spike-in can avoid most of these problems. Therefore, some studies also try to use stably expressed endogenous genes that can serve as internal references both in bulk (6) and single cell RNA-Seq (8,9). However, both of these two methods for scRNA-Seq perform a library size like normalization before detecting stable expressed genes, which automatically assume equal total RNA abundances and thus identify suboptimal stable genes when facing huge variations in total RNA abundances in heterogeneous single cell population. In addition, as scMerge is designed to handle and merge multiple batches, it does not suit cases when input dataset is from just one single batch. Another simple alternative to internal reference calculated the size factors for normalization based on just highly expressed genes (10).

Here, we develop an algorithm, ISnorm (Internal Spike-in-like-genes normalization) that selects a set of stably expressed genes (IS genes) as internal references and then to normalize scRNA-seq data accordingly. Notably, our algorithm selects genes based on the pairwise variance (a modified version of log ratio variance) (2) between IS genes from the input expression matrix and does not require any prior knowledge or the guidance of external reference datasets. We adopt this approach as previous work demonstrated that logratio variance based measurements of pairwise similarities outperformed Pearson’s correlation for compositional data such as RNA-Seq (11). In this work, we first demonstrate that ISnorm correctly selects a set of constantly expressed genes and provide unbiased estimate of size factors on simulated datasets. By applying ISnorm to several case study datasets, we also demonstrate that ISnorm improves the accuracy and enhances the statistical power of downstream analyses especially when transcriptome undergoes drastic changes.

## MATERIALS AND METHODS

### Overview ISnorm method

Here we gave a brief description of ISnorm algorithm (Figure 1). ISnorm first learned the internal variance for a set of high quality genes from raw matrix by calculating a slightly modified log- ratio variance (*LRV*) between the expression vectors of genes (2), called Dispersion of Ratio (*DoR*). Then ISnorm applied Density-Based Spatial Clustering of Applications with Noise (DBscan) algorithm to the distance matrix and selected several candidate genesets. For each candidate geneset, ISnorm calculate a statistical term called instability score for each single cell, reflecting the inconsistency of expression for these genes against a reference sample. Aggregation of the instability scores across all cells gave the reliability of the geneset. Finally, we selected the optimal geneset and used them for normalization.

**Figure 1.**
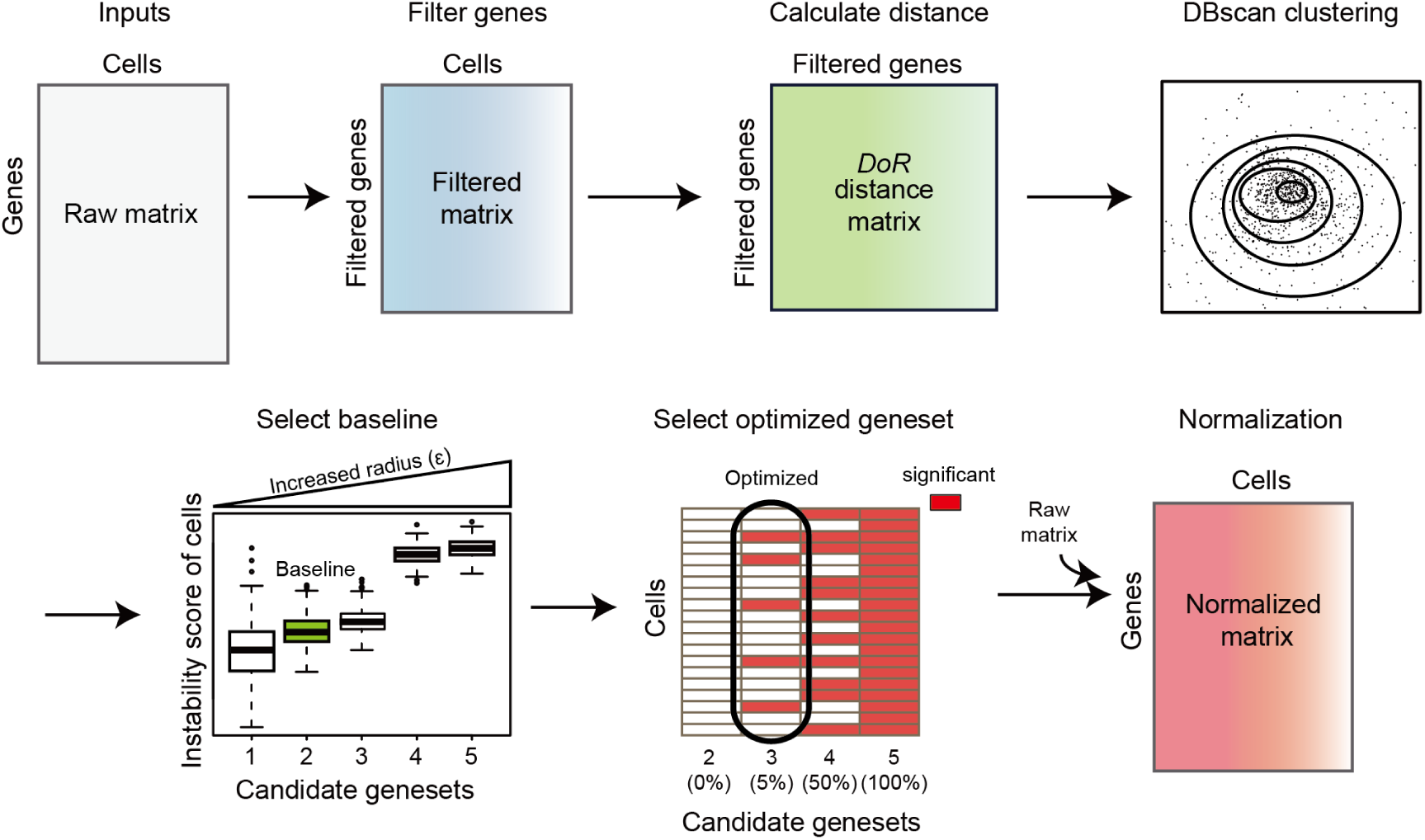
Overview of ISnorm pipeline.

#### Selection of a series of candidate genesets

ISnorm takes an expression matrix as inputs. Selecting a set of IS genes from a gene pool is the first step of ISnorm. As ISnorm simply ignores zero counts in normalization, it may perform poorly on genes with many zeros. Thus genes with less than 90% (default value) cells having nonzero expression are discarded. Assuming we have *n* cells and two non-negative vectors of gene expression ***X*** = (*x*_1_, *x*_2_, *… …, x*_*n*_)^*T*^ and ***Y*** = (*y*_1_, *y*_2_, *… …, y*_*n*_)^*T*^, we calculate the log ratio vector between ***X*** and ***Y*** by *z*_*i*_ = log_2_*x*_*i*_ *−* log_2_*y*_*i*_, and define *DoR* as follows:

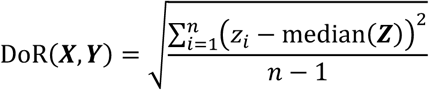

In ISnorm, *z*_*i*_ is not included if either *x*_*i*_ or *y*_*i*_ is zero. The only difference between *DoR* and squared root of *LRV* (2) is that we replaced the mean of ***Z*** by the median of ***Z***, because median values are more robust to outliers in single cell RNA-Seq due to undesirable variations. *DoR* is based on the assumption that the log ratio variance between two constantly expressed genes is always small (see Supplementary Note 1 for a detailed discussion on *DoR* and a brief comparison with its alternative solutions). Consequently, most constantly expressed genes (IS genes) will have small pairwise *DoR* values with each other, and form a region of high density in high dimensional space. To identify densely clustered IS genes, we apply DBscan algorithm in R package dbscan (12) on the pairwise distance matrix of *DoR*. Some tightly co-regulated genes may also have small *DoR* values with each other and cannot be easily distinguished from constantly expressed genes. Here we make the assumption that IS genes can outnumber these specific co-regulated genes in most cases and assign the genes in the largest cluster as IS genes. In DBscan clustering, the scanning radius (ε) and the minimum number of points (minPts, in this case it means minimum number of genes) need to be specified manually to identify a dense region but there is no golden rule to choose these two parameters. To overcome this problem, we feed a series of “expected number of IS genes” to ISnorm (the expected number starts from 5, gradually increases with a step of 5 and stops when it reaches 20% of the gene pool, throughout this study). ISnorm runs DBscan repeatedly with gradually increased ε and predicts several candidate IS genesets containing increasing number of IS genes. For simplicity, minPts is set to 5 in all cases (the default setting of R package dbscan).

#### Calculation of size factor based on geneset

For each candidate geneset, we calculate a size factor for each cell by the strategy provided by Lun *et al.* (13), with some modifications. Briefly, this strategy pools the information of all cells together to create a pseudo cell as reference and normalizes each cell against it. Assuming we have *n* cells and one candidate geneset with *m* IS genes, the reference is defined as follows:

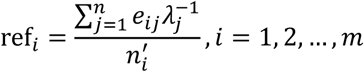

where ref_*i*_ denotes the reference expression for gene *i, e*_*ij*_ denotes the expression level of gene *i* in cell *j, λ*_*j*_ denotes a cell-specific coefficient adjusting the weight of each cell, which is the total counts of *m* IS genes in cell *j*, divided by the median value of total counts of *m* IS genes across all cells, and 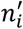 denotes the number of cells having nonzero expression for gene *i*. Therefore, the reference can be considered as weighted average expression levels for all IS genes across the whole cell population. By using 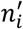 instead of *n*, we only consider nonzero expression when creating the reference. The size factor for cell *j* is calculated as follows (we only consider nonzero values here):

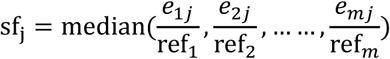

The size factor for each cell can be regarded as a median estimate of actual (this individual cell) versus averaged expression (from all cells) for all IS genes.

#### Selection of optimal geneset as IS genes

Finally we need to select an optimized candidate geneset best representing a set of true IS genes. If IS genes are correctly selected, we expect that the reference expression should have a strong linear relationship with the expression of IS genes in each cell. We define instability score for cell *j* as follows:

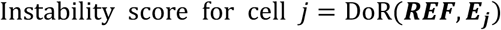

where ***REF*** = (ref_1_, ref_2_, *… …*, ref_*m*_)^*T*^ and ***E***_***j***_ = (*e*_1*j*_, *e*_2*j*_, *… …, e*_*mj*_)^*T*^. Unlike the *DoR* used to evaluate the similarity between two genes above, the *DoR* in instability scores is used to evaluate the similarities between a pseudo reference cell and any individual cells present in real single cell RNA-Seq data. Generally, candidate sets of larger size will result in higher instability scores for most cells, as more genes inconsistent with others are included. So a candidate set of smaller size and lower instability scores normally contains IS genes with higher confidence. But including slightly more IS genes can reduce the noises and lead to more reliable results. Thus we develop a strategy to select an optimal candidate set while constraining the instability scores of most cells to a reasonably low level. First we calculate the average instability score of all cells for each candidate geneset and select the largest candidate geneset with average instability score below a defined threshold. This candidate geneset is used as baseline to measure the intrinsic variation of dataset (the baseline geneset). We set the threshold to 0.1, empirically drawn from the real datasets investigated in this study. For some datasets of high heterogeneity or noises, the average instability score of all candidate sets may be larger than 0.1. In these cases we select the candidate set with lowest average instability score as the baseline geneset. Next we test whether larger genesets with more IS genes are still reasonably low on instability scores compared to the baseline geneset. As the instability score shares a close form of definition with *SD*, the instability scores of baseline geneset and an alternative candidate geneset are compared for each cell by F-test, with degrees of freedom to be the number of non-zero genes in baseline geneset and alternative geneset minus one. One-tailed *p* value is calculated to measure whether the instability score of geneset with more IS genes is larger than the one of the baseline geneset. The alternative geneset fails to pass the test if the percentage of cells showing statistical significance by F-test exceeds the imposed level (e.g. more than 5% cells with *p* value smaller than 0.05). The largest candidate geneset that meets the statistical threshold is selected as the optimal geneset. If no candidate geneset meets the threshold, the baseline geneset is selected.

Specifically, we found few mitochondrial genes and some external spike-ins (if added in experiment) may have extremely low instability scores and would always be reported. As relative abundances of mitochondrial genes increase significantly when cells undergo apoptosis, we removed them along with any external spike-ins before calculating *DoR* (see the second paragraph in Discussion for details).

### RNA-seq data processing

In this paper, we used several public single cell RNA sequencing data. For datasets with raw counts, we used processed raw counts from online database. For datasets lacking raw counts, we implemented a common pipeline to obtain raw gene expression count matrices. Raw reads were first processed by Trim Galore (Babraham Institute) to remove adapters and then aligned to *M. musculus* transcriptome (Ensembl v.38.89) or *H. sapiens* transcriptome (Ensembl v.37.87) merged with ERCC sequences if added in experiment using Bowtie v1.2.2 (14) or Bowtie2 v2.3.4.1 (15), depending on the read length. Raw counts and TPM values were estimated by RSEM-1.3.0 (16). Cells with fewer than 2,000 detected genes were filtered out, in addition to Bach dataset and human PBMCs dataset. We applied a more relaxed cell filtering step on Bach dataset and human PBMCs dataset (filtering cells with fewer than 1,000 detected genes) to test the computational efficiency of ISnorm. The effective library size was estimated by the summed expression of all genes after normalization upon different methods. The absolute mRNA content was defined as the ratio between summed counts of endogenous mRNA and ERCC spike-ins. For comparison, the median value of normalized or absolute mRNA content across all cells was scaled to the same value.

### Implementation of ISnorm and existing methods

Implementations of ISnorm, existing normalization methods and downstream analyses were carried out in R 3.4.3 unless otherwise noted. In DESeq2, Scran and SCnorm, default parameters were used when estimating size factors. Notably, user can specify the condition of each cell in Scran and SCnorm to avoid normalizing a highly heterogeneous cell population. But we did not specify it as there was no golden rule on how to set it. For library size normalization, size factors were defined as total counts divided by the median value of total counts of all cells.

### Performance comparison on simulation dataset

We compared the performance of ISnorm with two bulk-based methods, DESeq2 (17) and library size normalization, and two single-cell methods, Scran (13) and SCnorm (18), on the simulated datasets. All simulations were carried out in R package Splatter (19). Splatter estimates hyper parameters from real data to set most basic characteristics of simulated data. We estimated hyper parameters from one scRNA-seq data of mouse embryonic stem cells (mESCs) by Smart-seq2 protocol (20) and one scRNA-seq data of K562 cells by inDrop protocol (21) and simulated two datasets based on these parameters (Smart-seq2 simulation and inDrop simulation). Dropout option was turned on in Smart-seq2 simulation and turned off in inDrop simulation to match the sparsity of real data. In Smart-seq2 simulation, we also manually set common dispersion of biological coefficient of variation (BCV) from 0.46 to 0.23 to match the variation of real data. In each of two simulations, we simulated two subpopulations, each having 100 cells. Thirty percent of the genes were differentially expressed. The log fold-change of DE genes was sampled from a normal distribution (*μ* = 1, *σ* = 0.4). True size factors were defined as the library size multiplied by the sum of cell mean of all genes in the Splatter pipeline. Other simulation parameters were set to default values if unmentioned.

### Performance comparison on real datasets

To further evaluate the performance of ISnorm, we used five dataset, mouse preimplantation embryos (22), mouse embryogenesis (23), human embryogenesis (24), mouse preimplantation embryos ATAC-seq data (25), and glioblastoma (26), to evaluate the impact of different normalization methods on real datasets downstream analysis. Details for the five datasets analysis are described in Supplementary Methods.

### Bulk RNA-seq of CD3+ T cells and CD14+ monocytes

We conducted a bulk RNA-seq assay to validate the mRNA content difference between T cells and monocytes. PBMCs were isolated from healthy donors by Histopaque-1077 (Sigma#10771) according to the manufacturer instruction. Sorting 500,000 T lymphocytes and monocytes with BD FACS AriaIII indicated by specific maker (Supplemental Fig. S11), human CD3 (Biolegend#300440) and human CD14 (Biolegend#325620). Total RNA were isolated by TRIzol Reagent (Invitrogen#15596026), 1ul 10X diluted ERCC RNA spike-In Control Mixes (Invitrogen#4456740) were added to each sample for the cDNA library generation, sequencing library construction and data analysis. Three replicates were conducted for T lymphocytes and monocytes, separately.

### High Variable Gene (HVG) detection and motif enrichment analysis on PBMCs and K562 cells

Highly variable gene (HVG) detection is important to understand the heterogeneity of cell population and may also be affected by normalization. Here we showed that ISnorm improved performance of HVG detection compared to other normalization methods in human peripheral blood mononuclear cells (PBMCs). To get orthogonal evidence for benchmarking, we used the appearance of TF motifs with highly variable activity around the transcription starting sites (TSSs) of genes to measure their variation of expression. (Figure 5A). Although for one specific gene, it is always unreliable to predict its expression from epigenetic profiles, we considered that for a group of genes, the extent of enrichment of TF motifs with variable activity near the TSSs could reflect the variation of genes on average. HVGs inferred by better normalization methods were expected to have strong associations with variable TF motifs. In order to get a list of highly variable TF motifs, we applied chromVAR (22) to a single-cell ATAC-seq (scATAC-seq) dataset of human PBMCs and obtained a set of TF motifs that had the largest variation of chromatin accessibility across all cells. As the variations of single peaks cannot be inferred from nearly binary scATAC-seq data, chromVAR aggregated the reads from peaks containing the same motif to measure the activity level and variation of corresponding TF motif. After obtaining variable motifs, we estimated the enrichment of motifs by calculating the percentage of ATAC peaks containing a specific motif around the TSS of inferred HVGs upon ISnorm and other normalization methods. For each motif, the enrichment was measured by comparing frequency of its appearance in HVG associated peaks with the frequency in all peaks near TSS regions.

**Figure 2.**
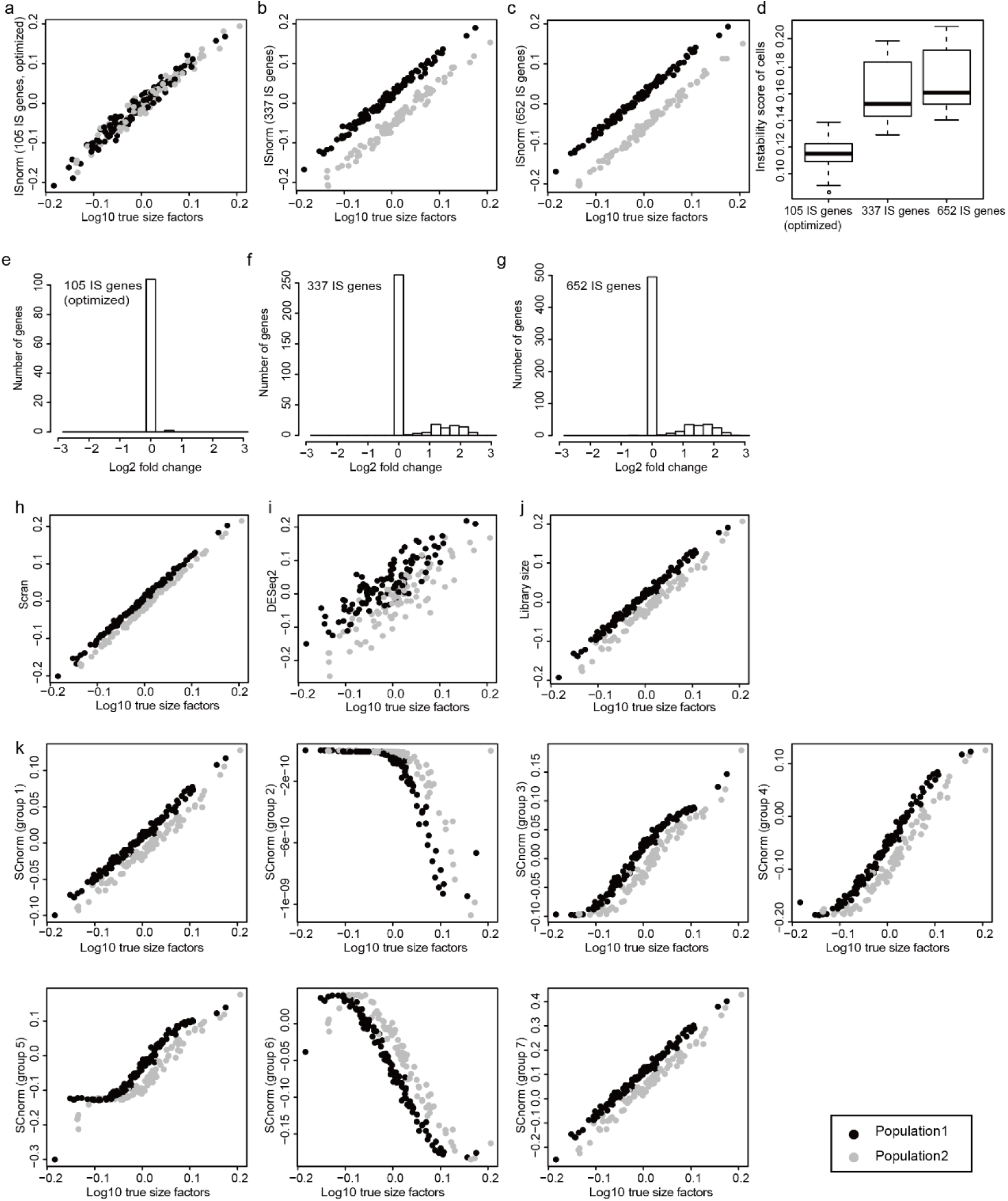
Performance of ISnorm and other existing methods on Smart-seq2 simulation. **a**-**c** Size factors estimated by ISnorm. **d** Instability scores of cells for each candidate set. **e**-**g** Distribution of log2 fold change of IS genes in each candidate set. **h**-**k** Size factors estimated by other existing methods. Black and grey dots represent cell population 1 and cell population 2, where 30% genes in population 2 are differentially expressed.

**Figure 3.**
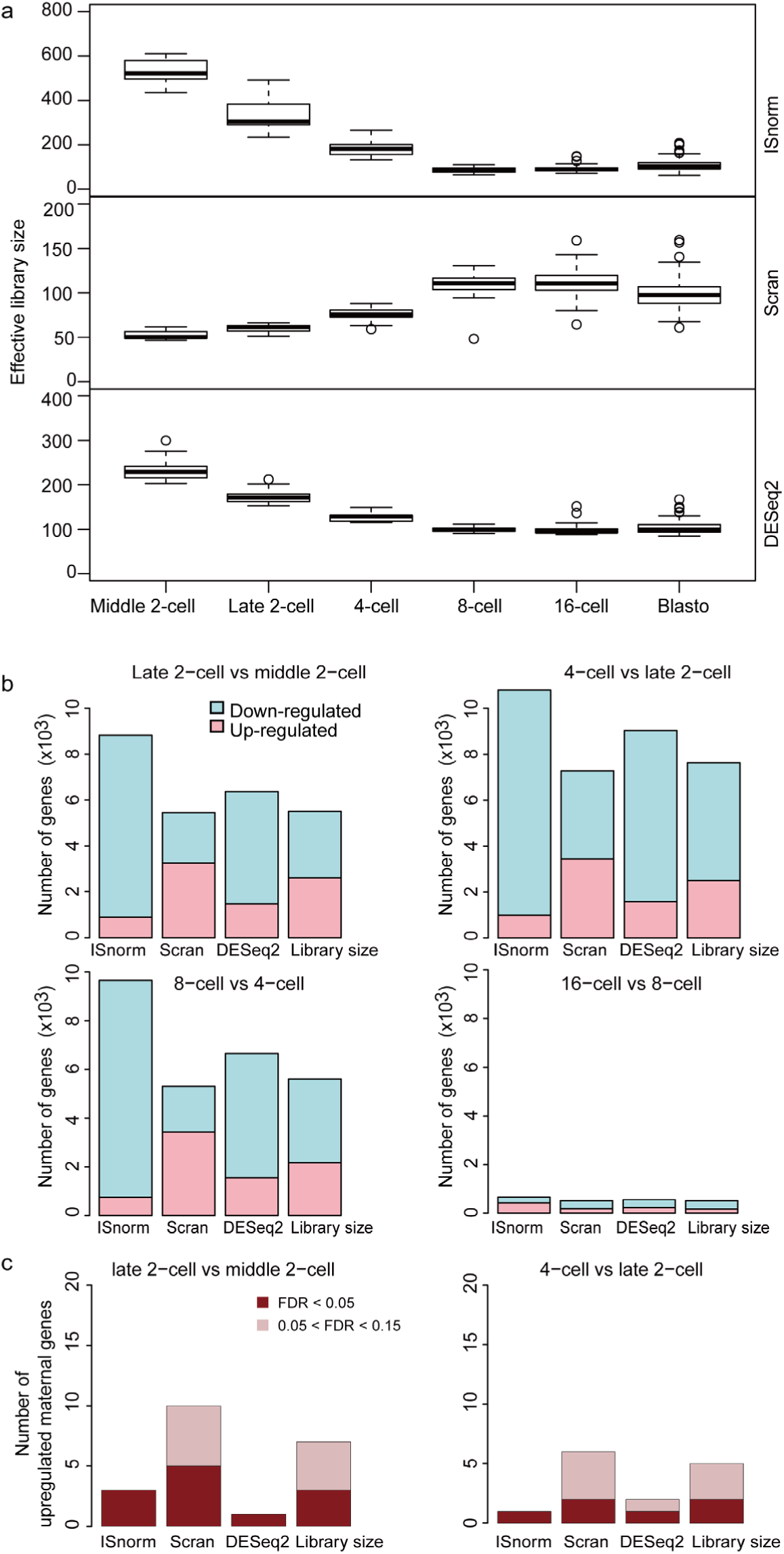
ISnorm detects drastic transcriptomic changes on Deng dataset of embryogenesis. **a** Effective library size estimated by ISnorm, Scran and DEseq2. **b** Number of genes detected as differentially expressed genes using DESeq2 by feeding size factors from ISnorm and other existing methods. **c** Number of maternal genes with log2 fold change > 0 and 0.05 < FDR < 0.15 (light red) and number of maternal genes with log2 fold change > 0 and FDR < 0.05 (dark red) inferred by feeding DESeq2 with size factors from ISnorm and other existing methods.

**Figure 4.**
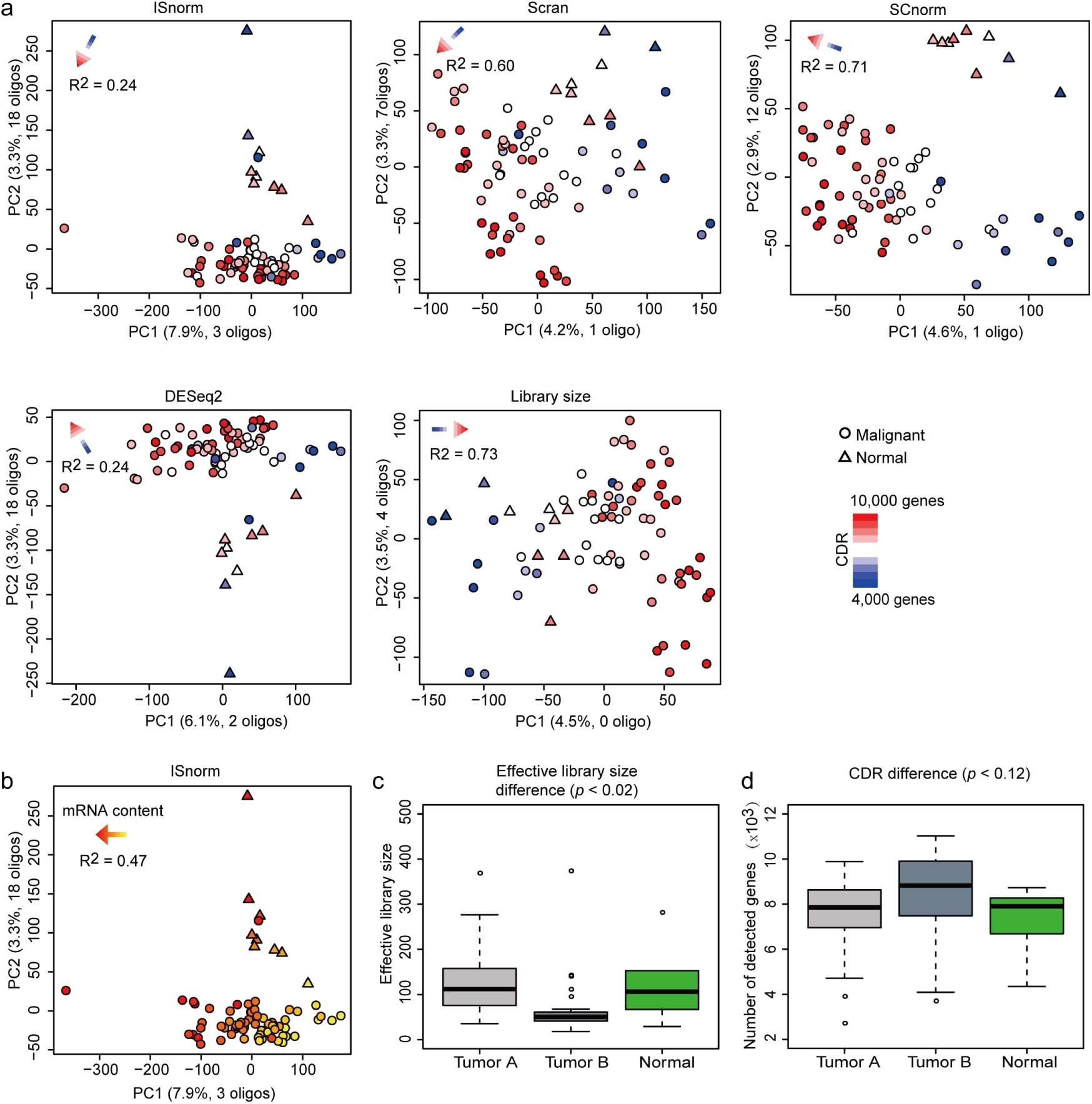
ISnorm improves the performance of PCA on Patel dataset. **a** The results of first two PCs inferred by PCA on normalized matrices. Malignant cells (circle) and normal cells (triangle) are colored according to the number of detected genes in each cell. The results of linear regression (CDR ∼ PC1 + PC2) are shown by an arrow on topleft. The separation of malignant and normal cells is more obvious in results based on ISnorm, SCnorm and DESeq2. **b** The same results of PCA based on ISnorm. Cells are colored according to log transformed effective library size estimated by ISnorm. **c-d** Comparison of **c** effective library size calculated by ISnorm and **d** number of detected genes for normal cells and two tumor clones. The *p* value is calculated by ANOVA.

**Figure 5.**
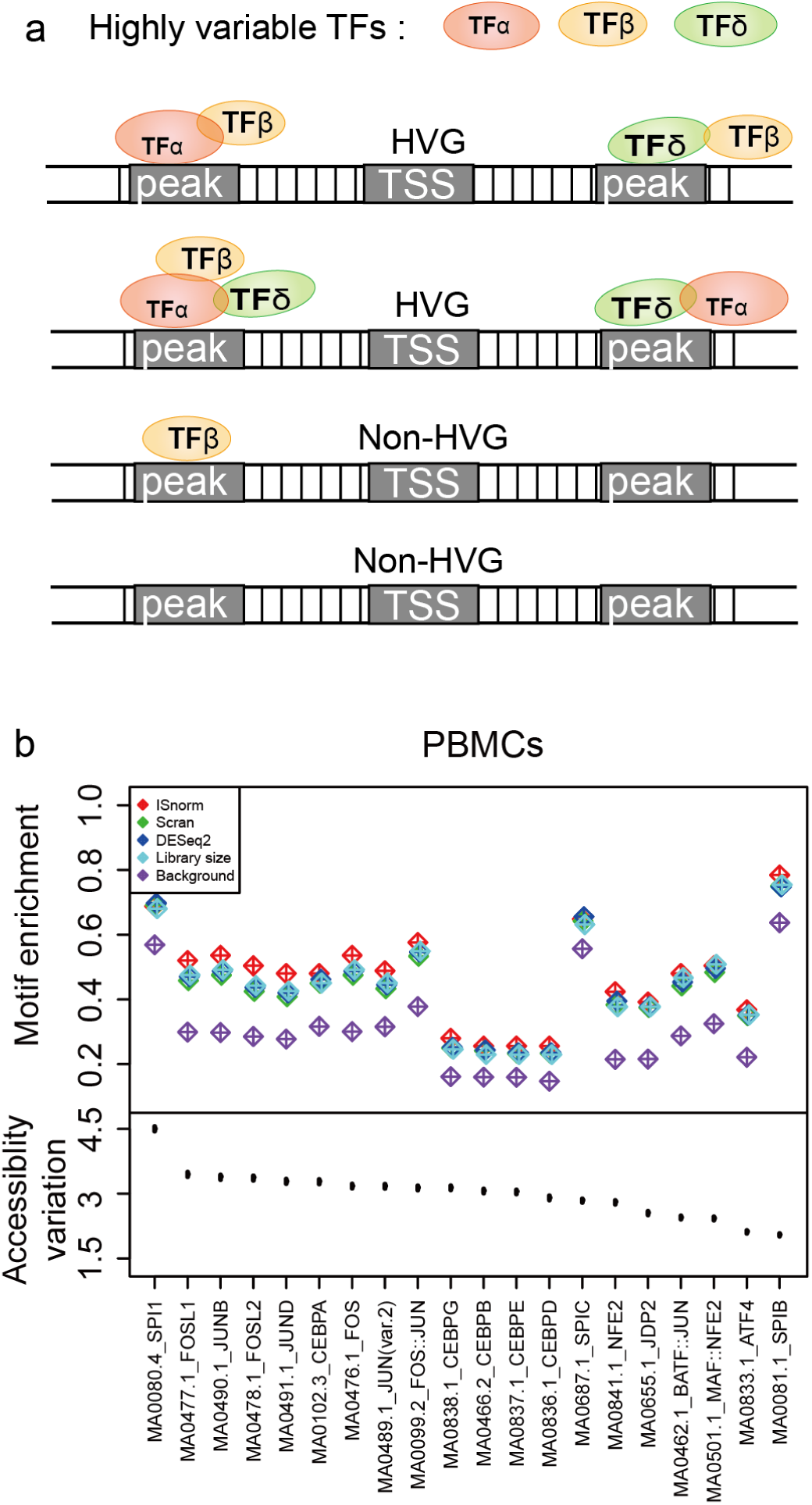
ISnorm improves performance of HVG detection in PBMCs dataset. **a** The schematic view of motif enrichment analysis. TFs with highly variable activity should be enriched in the regulatory elements of HVGs compared to non-HVGs. **b** The motif enrichment patterns of top 20 TF motifs with highest variability upon ISnorm and other normalization methods. The upper part shows the percentage of HVG associated peaks containing specific motif based on ISnorm and other normalization methods and the lower part shows the point estimates and error bars of accessibility variation of motifs from scATAC data.

The processed UMI count matrix of scRNA-seq dataset and count matrix of scATAC dataset of human PBMCs were downloaded from 10X Genomics Chromium website. We divided cells into CD3+ T cells, CD14+ monocytes and CD19+ B cells following the result of K-means clustering provided by 10X Genomics Chromium and manually checked it through the expression or the accessibility of promoter of several cell type specific markers. SCnorm failed to normalize the matrix within 24 hours and was not reported. For HVG detection on scRNA-seq datasets, we first applied a pre-filtering step to remove genes that were detectable in less than 50% cells, considering that dropouts could be a major source of variation for these genes. HVG detection was conducted using a decomposition method from R package scran (13) (trendVar and decomposeVar functions), which fitted a mean-variance trend to the normalized log-expression values of all endogenous genes and subtracted the technical variance from total variance. Given that there are only ∼800 genes left after pre-filtering step, the top 100 genes with lowest FDR values were selected as HVGs.

In scATAC datasets, peaks regions were obtained from 10X Genomics Chromium website for PBMCs or called by MACS2 (27) on merged data of single cells for K562 cells. Peaks that were detectable in more than 10% cells in scATAC dataset and within 10kb upstream and downstream regions of the TSS sites of HVGs were considered as associated peaks with HVGs detected in previous steps. The top 20 TF motifs with highest variances calculated through chromVAR (28) from scATAC dataset were selected as highly variable TF motifs. The location of motifs was obtained through R package JASPAR2016. Then we examined whether these highly variable TF motifs were more likely to appear in HVG associated peaks, which gave the evaluation of HVG detection on scRNA-seq data. The enrichments of other motifs were similar for all normalization methods and thus were not shown.

To confirm that the motif enrichment analysis described above did not introduce any overfitting problems, we also analyzed single cell data of K562 cells that were supposed to be homogeneous. As expected, ISnorm performed similarly with other normalization methods and the enrichment patterns were globally weak for all TFs (Supplemental Fig. S7D).

## RESULTS

### ISnorm correctly estimates true size factors in simulated datasets

We first compared the performance of ISnorm with two bulk-based methods, DESeq2 (17) and library size normalization, and two single-cell methods, Scran (13) and SCnorm (18), on the simulated datasets. We conducted two simulations in R package Splatter (19) to mimic real data from two scRNA-seq protocols, Smart-seq2 (29) and inDrop (21), respectively (Smart-seq2 simulation and inDrop simulation). Each simulated dataset contained two subpopulations and 30% DE genes between two subpopulations, which represented a typical strong DE case. SCnorm failed to calculate a normalized matrix in inDrop simulation and was not reported. This may be due to the fact that SCnorm is not designed to handle data with high sparsity as described in its manual. We found that ISnorm provided the best estimate of true size factors and most genes selected by ISnorm were constantly expressed (non-DE) across all cells when constraining the instability score of most cells in low level (Figures 2A-2G and Supplemental Fig. S1A-S1G). In Smart-seq2 simulation, candidate genesets with 337 and 652 IS genes contained large proportion of DE genes (Figures 2F and 2G) and led to biased results similar to other methods (Figures 2B and 2C). However, these two candidate genesets also showed significantly higher instability scores compared to the optimal geneset with 105 IS genes (Figure 2D), which suggested that our method was quite effective in excluding undesirable IS genes (such as DE genes). All other existing methods provided larger estimates of size factors for cells in the first subpopulation, exhibiting subpopulation-based biases (two populations clearly separate from each other on the scatter plots in Figures 2H-2K and Supplemental Fig. S1H-S1J).

### ISnorm can detect drastic changes of cellular mRNA abundance in a mixed population

To further evaluate the performance of ISnorm, we examined the impact of different normalization methods on several real datasets. The first dataset contains single cell RNA-seq data of mouse preimplantation embryos from Deng *et al.* (22), a developmental process with drastic transcriptome changes. Normalized expression matrices were calculated by ISnorm and other normalization methods. We only analyzed cells from middle 2-cell to blastocyst stage as we found cells from zygote or early 2-cell stages showed significantly higher instability scores and thus didn’t share common IS genes with other cells (Supplemental Fig. S2). We first estimated the effective library size (mRNA content after normalization) to show the overall trend of differential expression. ISnorm and DESeq2 suggested a clear decrease in effective library size from middle 2-cell to 8-cell stage (Figure 3A). The decrease of mRNA content through early embryogenesis has been demonstrated by a scRNA-seq data with external spike-ins (23) (Supplemental Fig. S3). Scran normalization failed to reveal this pattern. We next evaluated the impact of different normalization methods on DE gene detection using DESeq2 (17). Differential expression analysis based on ISnorm detected more down-regulated genes than other methods from middle 2-cell to 8- cell stages, which was consistent with the decrease in mRNA content (Figure 3B). It is usually difficult to decide a gene to be up- or down-regulated by computational methods but it is known that genes with a silenced epigenetic state are less likely to be up-regulated. Thus we selected 665 genes with high expression but closed chromatin state in 2-cell and 4-cell stages (these genes are usually referred to as maternal genes that go through maternal clearance during embryogenesis) by the combined information of scRNA-seq data (22) and ATAC-seq data (25) of mouse preimplantation embryos (Supplemental Fig. S4) for benchmarking. As expected, ISnorm controlled the number of up-regulated maternal genes. DESeq2 also showed similar results with ISnorm while other methods always suggested more maternal genes to be up-regulated (Figure 3C).

We also tried ISnorm on several other studies to examine whether ISnorm could reliably detect cellular mRNA content differences due to strong differential expression. ISnorm generated similar decreased patterns in effective library size for another dataset of mouse embryogenesis (23) and one dataset of human embryogenesis (24), while other methods including DESeq2 failed to do so (Supplemental Fig. S5A-S5B). In human peripheral blood mononuclear cells (PBMCs), ISnorm suggested higher mRNA content of CD14+ monocytes than CD3+ T cells, agreeing with total UMI counts of single cells, which was a precise estimate of mRNA content when samples were sequenced to saturation (7). We also validated this result through a bulk RNA-seq assay with External RNA Controls Consortium (ERCC) spike-ins (Supplemental Fig. S5C; see Methods for details). In mouse hematopoietic stem and progenitor cells (HSPCs) (30), We found that an unexpected pattern of heterogeneity, where few cells have significantly larger library size compared to their size factors estimated by ISnorm, suggesting that they might have higher mRNA content than the majority of cells (Supplemental Fig. S5D). The increased mRNA content of these outlier cells could also be validated by ERCC spike-ins.

Previous study indicates that cells in G2/M stages have higher mRNA content than G1 and S stages (7). However, ISnorm did not detect higher mRNA content for cells in G2/M stages in Leng dataset (31) (Supplemental Fig. S5E). We argue that this is because ISnorm accounts for mRNA content difference due to partial transcriptome change (as far as the IS genes are not affected) but normalizes mRNA content difference due to whole transcriptome change (e.g. cellular volume increase in G2/M stage). Thus, unlike external spike-ins, no further normalization is required to account for the difference in cellular volumes, which affect the global transcriptional output (32). This also helps explain the fact that ISnorm could not fully capture the change of mRNA content estimated by external spike-ins for embryogenesis (Supplemental Fig. S3).

### ISnorm improves the performance of downstream analysis in a case study of tumor tissue

As cells in cancer tissues show much stronger heterogeneity, we wonder whether ISnorm performs well for complex cell populations such as cancer. To address this question, we evaluated the performance of ISnorm on a case study of glioblastoma (26). We only showed results from patient MGH31 as it contained two tumor clones and non-malignant cells (Supplemental Fig. S6A), which can serve as ground truth. We found that PCA upon ISnorm, DESeq2 and SCnorm roughly distinguished malignant cells from normal cells by PC1 and PC2, while Scran and library size normalization failed to do so (Figure 4A). Instead, the number of detected genes (cellular detection rate, CDR) of single cells was a dominant factor revealed by PCA upon Scran and library size normalization (*R*^2^ >= 0.6; CDR was also a dominant factor in PCA upon SCnorm but did not affect the classification of cells), suggesting the first two PCs based on these normalization methods may mainly capture the technical signals. But all normalization methods successfully distinguished malignant cells from normal cells by three PCs (Supplemental Fig. S6B). Previous study suggested variation in CDR can easily overwhelm true biological information and should be controlled (33). As ISnorm controls the technical noises better, we assume that genes contributing to PC1 and PC2 may capture more biologically relevant genes. Further investigation on genes with high loadings in PC1 or PC2 upon ISnorm showed that 21 of them overlapped with the known oligodendrocyte markers from the MSigDB database (Figure 4A). Other methods showed similar pattern but with fewer overlapped genes. Moreover, we found that PC1 upon ISnorm explained more variations than other methods (Figure 4A) and mainly reflected the effective library size of tumor cells estimated by ISnorm (*R*^2^ = 0.47; Figure 4B). Coincidently, cells from one tumor clone had significantly lower levels of effective library size compared with other cells (Figure 4C). We also tested the variation of CDR between three types of cells but found no significant difference (Figure 4D). These results suggested that PC1 might capture the differences in cellular RNA abundance among tumor cells. Lacking external spike-ins, we could not validate the mRNA content estimated by ISnorm. But we found that it was highly consistent with library size of single cells (Supplemental Fig. S6C), which might reflect the amount of starting mRNA materials, thus the mRNA content for each cell (see Supplementary Note 2 for more details). These facts together indicated that PCA upon ISnorm captures the true biological variation rather than technical noises. Additionally, we found DESeq2 performed similarly as ISnorm in this case study, except for less variation explained than ISnorm on PC1.

### ISnorm improves performance of HVG detection in human PBMCs

In PBMCs, we found that motifs with variable activity showed significant enrichment patterns around the TSSs of HVGs upon all normalization methods, proving that our hypothesis on the enrichment of variable motifs around TSSs of HVGs was true (34). Among all normalization methods, HVGs upon ISnorm showed the most significant enrichment patterns for motifs with high variations (Figure 5B), suggesting that the HVGs upon ISnorm agreed better with epigenetic profiling of single cells. We also divided the PBMCs into three more homogeneous populations, CD3+ T cells, CD14+ monocytes and CD19+ B cells and applied the motif enrichment analysis to test whether ISnorm still performed better. We found that ISnorm showed improvements for some of the motifs for T cells but performed equally for monocytes and B cells compared to other methods (Supplemental Fig. S7A-S7C). These findings also agreed with the fact that ISnorm showed strong confliction with library size for PBMCs but less inconsistency for T cells, monocytes and B cells separately (Pearson Correlation Coefficient of size factors between ISnorm and library size: PBMCs, 0.69; T cells, 0.79; monocytes, 0.93; B cells, 0.84).

### ISnorm identifies cell-specific constantly expressed genes

One advantage of ISnorm is that it can predict a set of constantly expressed genes as IS genes. We applied ISnorm to different datasets and multiple tissues in the same dataset to examine generalizability of selected IS genes (Supplemental Table S2 and Supplementary Table S3). Although in most cases housekeeping genes were selected as IS genes, we also found a set of tissue-specific IS genes (e.g. in brain, kidney, liver epithelial cells; Supplemental Table S3). These findings may help researchers discover population-specific constantly expressed genes that can be used in other applications such as single-cell qPCR. However, we also noted that for the dataset containing diverse cell types, ISnorm may fail due to the fact that there are no common IS genes shared by all cells.

### The computation time and resource requirements of ISnorm

To confirm that ISnorm is generally applicable to large droplet-based datasets, we test the computational efficiency of ISnorm on two datasets with more than 10,000 cells generated by 10X Genomics Chromium platform, one dataset of mammary epithelial cells from Bach *et al.* (35) (∼20,000 cells) and PBMCs dataset (∼10,000 cells). As the running time of ISnorm depends on the number of input genes as well as the number of cells, we tried two gene inputs (genes having nonzero expression in more than 90% and 70% cells). The reported IS genes were identical from two gene inputs (Supplemental Table S2). The running time of ISnorm was always within minutes. Detailed information on the size of datasets and running time was shown in Supplemental Table S4. Analyses were performed on an I840-G25 computation workstation with Intel(R) Xeon(R) E7-8880v3 processors, under one thread and 8GB RAM limitations.

## DISCUSSION

Through benchmarking on simulated datasets, we found that all existing normalization methods showed subpopulation-specific biases under strong DE case. But in Deng dataset of embryogenesis and Patel dataset of glioblastoma, the results upon DESeq2 normalization were similar with ISnorm and were better compared to other methods. Previous study suggested that high sparsity of the data had a strong effect on DESeq2, as the calculation of the pseudo sample which served as reference was only well defined on a limited number of genes that were detected in all cells, which might be problematic (7). However, in scRNA-seq data, genes with high expression level are less affected by dropouts and with lower Coefficients of Variation (CV) (1), which may provide more reliable estimate on the size factor of the cell than low expressed genes. Actually, our work suggested that a small set of constantly expressed genes was enough for robust normalization. Thus the improved performance of DESeq2 on the case study datasets mentioned above can be explained by the usage of non-zero count genes, which also happen to exhibit features of IS genes. However, by applying existing normalization methods on several datasets of embryogenesis, we found that the non-zero count genes used in DESeq2 normalization did not always behave as IS genes, and failed to reflect the true patterns of transcriptomic changes. Thus the performance of DESeq2 relies on the percentage of IS genes used to calculate the reference and may vary significantly among different datasets.

ISnorm selects a set of constantly expressed genes to normalize the data. But we found that normally there was no clear boundary between constantly expressed genes and differentially expressed genes in real datasets. Thus the strategy for identifying IS genes is based on researchers’ preferences: fewer genes with lower variance or more genes with higher variance. In most cases, we found different selecting strategies did not affect the results much as the size factors estimated by different candidate genesets were highly consistent given they always shared many common genes (Supplemental Fig. S8 and S9). However, we did find that in some datasets different candidate genesets might share no common genes and generated fairly different results (e.g. P0205 and P0508 in Zheng dataset, Supplemental Fig. S9; Leng dataset, Supplemental Fig. S10). For P0205 and P0508 in Zheng dataset, we thought the optimal geneset containing *HLA* genes was more likely to be true IS genes given the fact that ISnorm reported the same genes in *HLA* family in other three patients (Supplemental Fig. S9; Supplemental Table S2. Additionally, the instability scores of two non-optimal genesets were considerably higher and showed strong deviance from an empirical null distribution (see Supplementary Note 3 for details). But given such few IS genes, the results of ISnorm did not prove to be reliable. For Leng dataset, five mitochondrial genes were reported and showed inconsistency with other nuclear candidate genesets (Supplemental Fig. S10). There is no surprise that some mitochondrial genes may have low internal variance with each other given their important biological functions. As mitochondrial genes are different from nuclear genes in transcription, they might easily lead to gene-specific bias. Unlike P0205 and P0508 in Zheng dataset, we found that the instability scores of other nuclear candidate genesets with much more genes were still low enough to be considered as good IS genes (Supplemental Fig. S10). Given these results, we thought that instability scores proved to be a good indicator of true IS genes and constraining the instability scores of IS genes in a reasonably low level could prevent both gene-specific bias and selection of highly variable genes. However, if the optimal geneset contained few genes and showed inconsistency with other non-optimal genesets, it might mean that there were few or no IS genes shared by cells. In these cases it is safer to choose the geneset with lower instability scores but the results should be treated carefully.

Through benchmarking in this study, we found that in most cases all normalization methods provided similar estimates of size factors except for ISnorm. This is no surprise as they all share similar underlying assumptions that the majority of transcriptome remains constant. But with a different underlying assumption that only IS genes need to be constantly expressed, ISnorm showed strong confliction with other methods for some datasets. We then consider that ISnorm is not just a technical improvement in normalization but changes the basic concept of how to choose the baseline for gene expression measurement. We found that such difference in assumption would mainly affect the results of DE analysis, but it is hard to directly decide which assumption is more appropriate. However, we demonstrated that ISnorm showed higher consistency with epigenetic profiling for heterogeneous cell population (in mouse embryogenesis and human PBMCs dataset) and improved the performance PCA and subsequent informative gene extraction when the whole transcriptome underwent significant changes (Patel dataset of glioblastoma). These results implied that ISnorm would help reveal new biological patterns from single cell data.

From the results above, ISnorm proved to be most useful when the cellular total RNA abundances exhibited drastic variations among the cell population. When available, a UMI total count per cell in general is a good read out of cellular total RNA abundance. Otherwise, an ISnorm normalized total library size can be used. Thus, if the major clusters of cell populations showed significant differences in cellular total RNA abundances either in the form of UMI counts or ISnorm normalized library sizes (Figure 3A, Figure S5 C, D), ISnorm may give an alternative view on, and in many cases improve accuracies on the detection of, differentially expressed or highly variable genes.

Our results also demonstrated that the assumption of ISnorm might be violated in cases where distinct sets of IS genes were identified in different cell subpopulations. For example, mouse liver is mostly consisted of hepatocytes, but also a small number of non-hepatocyte cells. IS genesets identified in hepatocytes and non-hepatocytes cells separately did not overlap (10X protocol in Table S3). Because hepatocyte is the dominant cell type, IS genes identified for liver tissue (>95% hepatocytes) were consistent with IS genes for hepatocyte. Applying the same IS geneset learned from hepatocytes to non-hepatocytes didn’t fulfill the assumption of ISnorm, and thus would lead to incorrect size factor estimates for these cells (Fig. S12). In addition to ensure IS genesets identified in different subpopulation agree well, it is also crucial to evaluate the reliability of IS genes for any given cell cluster/population. Generally, the number of genes and the instability score of optimal geneset are good indicators of the reliability of ISnorm. If the number of IS genes is very low (e.g. < 10) or the instability score of optimal geneset is very high (e.g. > 0.2), ISnorm may easily fail given there are no good IS genes. Moreover, users should also be cautious against the cells will have significantly high instability scores (see the instability scores of zygotes and cells in early 2-cell stage in Fig. S2). Extreme high instability scores suggest IS genes identified in the whole population do not behavior similarly in these cells. Alternatively, extremely high instability scores can serve as an indicator that further separation of cell population is needed.

In summary, we introduced ISnorm to normalize scRNA-seq data based on a set of IS genes learned from input data. Our method to identify proper IS genes built upon a *DoR* (a modified version of *LRV* from compositional data analysis) calculation to evaluate distances between genes, suggesting that it is maybe worthwhile to explore the potential application of compositional data analysis methodology in the field of NGS analysis, especially for the noisy single cell data. Our approach is based on a relaxed assumption compared to existing methods that only a small number of genes are constantly expressed in a mixed cell population. Evaluation on simulated datasets indicated that ISnorm correctly identified constantly expressed genes and provided unbiased estimate of size factors in the case of strong DE. By applying ISnorm on several case study datasets, we demonstrated that our approach not only reveals the heterogeneity of gene expression and cellular mRNA abundance among individual cells, but also improves the performance of downstream analyses.

## Supporting information

no link

## DATA AVAILABILITY

All scRNA-seq datasets from published literatures or websites are summarized in Supplementary Table S1. All raw and processed sequencing data generated in this study have been submitted to the Genome Sequence Archive (GSA: https://bigd.big.ac.cn/gsa.) (36) in BIG Data Center (37), Beijing Institute of Genomics (BIG), Chinese Academy of Sciences, under temporary link https://bigd.big.ac.cn/gsa-human/s/TW1RvpE0. ISnorm is available under an MIT license on Github at https://github.com/bioliyezhang/ISnorm.

## SUPPLEMENTARY DATA

**Supplementary.pdf**

## ACKNOWLEDGEMENTS

We would like to thank the anonymous reviewers for their detailed comments and suggestions for the manuscript. We would like to thank Fangqing Zhao, Xiaoqi Zheng, Naomi Altman, Jie Zheng, Weituo Zhang, Lei Lei for useful comments and suggestions on the manuscript. We would like to thank Yingdong Zhang on his technical support on the HPC platform of ShanghaiTech University.

## REFERENCES

1. Brennecke, P., Anders, S., Kim, J.K., Kolodziejczyk, A.A., Zhang, X., Proserpio, V., Baying, B., Benes, V., Teichmann, S.A., Marioni, J.C. et al. (2013) Accounting for technical noise in single-cell RNA-seq experiments. Nature methods, 10, 1093–1095.

2. Aitchison, J. (1986) The Statistical Analysis of Compositional Data.

3. Quinn, T.P., Erb, I., Richardson, M.F. and Crowley, T.M. (2018) Understanding sequencing data as compositions: an outlook and review. Bioinformatics (Oxford, England), 34, 2870–2878.

4. Erb, I. and Notredame, C. (2016) How should we measure proportionality on relative gene expression data? Theory in biosciences = Theorie in den Biowissenschaften, 135, 21–36.

5. Jiang, L., Schlesinger, F., Davis, C.A., Zhang, Y., Li, R., Salit, M., Gingeras, T.R. and Oliver, B. (2011) Synthetic spike-in standards for RNA-seq experiments. Genome research, 21, 1543–1551.

6. Risso, D., Ngai, J., Speed, T.P. and Dudoit, S. (2014) Normalization of RNA-seq data using factor analysis of control genes or samples. Nature biotechnology, 32, 896–902.

7. Vallejos, C.A., Risso, D., Scialdone, A., Dudoit, S. and Marioni, J.C. (2017) Normalizing single-cell RNA sequencing data: challenges and opportunities. Nature methods, 14, 565–571.

8. Yip, S.H., Wang, P., Kocher, J.A., Sham, P.C. and Wang, J. (2017) Linnorm: improved statistical analysis for single cell RNA-seq expression data. Nucleic acids research, 45, e179.

9. Lin, Y., Ghazanfar, S., Wang, K.Y.X., Gagnon-Bartsch, J.A., Lo, K.K., Su, X., Han, Z.G., Ormerod, J.T., Speed, T.P., Yang, P. et al. (2019) scMerge leverages factor analysis, stable expression, and pseudoreplication to merge multiple single-cell RNA-seq datasets. Proceedings of the National Academy of Sciences of the United States of America, 116, 9775–9784.

10. Weinreb, C., Wolock, S. and Klein, A.M. (2017) SPRING: a kinetic interface for visualizing high dimensional single-cell expression data. Bioinformatics (Oxford, England), 34, 1246–1248.

11. Lovell, D., Pawlowsky-Glahn, V., Egozcue, J.J., Marguerat, S. and Bahler, J. (2015) Proportionality: a valid alternative to correlation for relative data. PLoS computational biology, 11, e1004075.

12. Ester, M., Kriegel, H.-P., Sander, J. and Xu, X. (1996) A density-based algorithm for discovering clusters in large spatial databases with noise. KDD, 226–231.

13. Lun, A.T., Bach, K. and Marioni, J.C. (2016) Pooling across cells to normalize single-cell RNA sequencing data with many zero counts. Genome Biol, 17, 75.

14. Langmead, B., Trapnell, C., Pop, M. and Salzberg, S.L. (2009) Ultrafast and memory-efficient alignment of short DNA sequences to the human genome. Genome Biol, 10, R25.

15. Langmead, B. and Salzberg, S.L. (2012) Fast gapped-read alignment with Bowtie 2. Nature methods, 9, 357–359.

16. Li, B. and Dewey, C.N. (2011) RSEM: accurate transcript quantification from RNA-Seq data with or without a reference genome. BMC bioinformatics, 12, 323.

17. Love, M.I., Huber, W. and Anders, S. (2014) Moderated estimation of fold change and dispersion for RNA-seq data with DESeq2. Genome Biol, 15, 550.

18. Bacher, R., Chu, L.F., Leng, N., Gasch, A.P., Thomson, J.A., Stewart, R.M., Newton, M. and Kendziorski, C. (2017) SCnorm: robust normalization of single-cell RNA-seq data. Nature methods, 14, 584–586.

19. Zappia, L., Phipson, B. and Oshlack, A. (2017) Splatter: simulation of single-cell RNA sequencing data. Genome Biol, 18, 174.

20. Ziegenhain, C., Vieth, B., Parekh, S., Reinius, B., Guillaumet-Adkins, A., Smets, M., Leonhardt, H., Heyn, H., Hellmann, I. and Enard, W. (2017) Comparative Analysis of Single-Cell RNA Sequencing Methods. Molecular cell, 65, 631-643.e634.

21. Klein, A.M., Mazutis, L., Akartuna, I., Tallapragada, N., Veres, A., Li, V., Peshkin, L., Weitz, D.A. and Kirschner, M.W. (2015) Droplet barcoding for single-cell transcriptomics applied to embryonic stem cells. Cell, 161, 1187–1201.

22. Deng, Q., Ramskold, D., Reinius, B. and Sandberg, R. (2014) Single-cell RNA-seq reveals dynamic, random monoallelic gene expression in mammalian cells. Science (New York, N.Y.), 343, 193–196.

23. Goolam, M., Scialdone, A., Graham, S.J.L., Macaulay, I.C., Jedrusik, A., Hupalowska, A., Voet, T., Marioni, J.C. and Zernicka-Goetz, M. (2016) Heterogeneity in Oct4 and Sox2 Targets Biases Cell Fate in 4-Cell Mouse Embryos. Cell, 165, 61–74.

24. Yan, L., Yang, M., Guo, H., Yang, L., Wu, J., Li, R., Liu, P., Lian, Y., Zheng, X., Yan, J. et al. (2013) Single-cell RNA-Seq profiling of human preimplantation embryos and embryonic stem cells. Nature structural & molecular biology, 20, 1131–1139.

25. Wu, J., Huang, B., Chen, H., Yin, Q., Liu, Y., Xiang, Y., Zhang, B., Liu, B., Wang, Q., Xia, W. et al. (2016) The landscape of accessible chromatin in mammalian preimplantation embryos. Nature, 534, 652–657.

26. Patel, A.P., Tirosh, I., Trombetta, J.J., Shalek, A.K., Gillespie, S.M., Wakimoto, H., Cahill, D.P., Nahed, B.V., Curry, W.T., Martuza, R.L. et al. (2014) Single-cell RNA-seq highlights intratumoral heterogeneity in primary glioblastoma. Science (New York, N.Y.), 344, 1396–1401.

27. Zhang, Y., Liu, T., Meyer, C.A., Eeckhoute, J., Johnson, D.S., Bernstein, B.E., Nusbaum, C., Myers, R.M., Brown, M., Li, W. et al. (2008) Model-based analysis of ChIP-Seq (MACS). Genome Biol, 9, R137.

28. Schep, A.N., Wu, B., Buenrostro, J.D. and Greenleaf, W.J. (2017) chromVAR: inferring transcription-factor-associated accessibility from single-cell epigenomic data. Nature methods, 14, 975–978.

29. Picelli, S., Faridani, O.R., Bjorklund, A.K., Winberg, G., Sagasser, S. and Sandberg, R. (2014) Full-length RNA-seq from single cells using Smart-seq2. Nature protocols, 9, 171–181.

30. Nestorowa, S., Hamey, F.K., Pijuan Sala, B., Diamanti, E., Shepherd, M., Laurenti, E., Wilson, N.K., Kent, D.G. and Gottgens, B. (2016) A single-cell resolution map of mouse hematopoietic stem and progenitor cell differentiation. Blood, 128, e20–31.

31. Leng, N., Chu, L.F., Barry, C., Li, Y., Choi, J., Li, X., Jiang, P., Stewart, R.M., Thomson, J.A. and Kendziorski, C. (2015) Oscope identifies oscillatory genes in unsynchronized single-cell RNA-seq experiments. Nature methods, 12, 947–950.

32. Padovan-Merhar, O., Nair, G.P., Biaesch, A.G., Mayer, A., Scarfone, S., Foley, S.W., Wu, A.R., Churchman, L.S., Singh, A. and Raj, A. (2015) Single mammalian cells compensate for differences in cellular volume and DNA copy number through independent global transcriptional mechanisms. Molecular cell, 58, 339–352.

33. Finak, G., McDavid, A., Yajima, M., Deng, J., Gersuk, V., Shalek, A.K., Slichter, C.K., Miller, H.W., McElrath, M.J., Prlic, M. et al. (2015) MAST: a flexible statistical framework for assessing transcriptional changes and characterizing heterogeneity in single-cell RNA sequencing data. Genome Biol, 16, 278.

34. Cao, J., Cusanovich, D.A., Ramani, V., Aghamirzaie, D., Pliner, H.A., Hill, A.J., Daza, R.M., McFaline-Figueroa, J.L., Packer, J.S., Christiansen, L. et al. (2018) Joint profiling of chromatin accessibility and gene expression in thousands of single cells. Science (New York, N.Y.), 361, 1380–1385.

35. Bach, K., Pensa, S., Grzelak, M., Hadfield, J., Adams, D.J., Marioni, J.C. and Khaled, W.T. (2017) Differentiation dynamics of mammary epithelial cells revealed by single-cell RNA sequencing. Nat Commun, 8, 2128.

36. Wang, Y., Song, F., Zhu, J., Zhang, S., Yang, Y., Chen, T., Tang, B., Dong, L., Ding, N., Zhang, Q. et al. (2017) GSA: Genome Sequence Archive. Genomics, proteomics & bioinformatics, 15, 14–18.

37. Members, B.D.C. (2019) Database Resources of the BIG Data Center in 2019. Nucleic acids research, 47, D8–d14.

