## Supplementary material for "Normalizing single-cell RNA sequencing data with internal spike-in-like genes": no link

### 1 Supplementary Methods

#### 1.1 Deng, Goolam and Yan datasets processing

Cells from zygote and early 2-cell stages in Deng dataset (1) showed higher instability scores than other cells (Supplemental Fig. S2), suggesting that identified IS genes may be differentially expressed in these two stages. The ATAC-seq data also indicated that the identified IS genes lacked strong signals in early 2-cell stages (Supplemental Fig. S4). Thus we excluded cells in zygote or early 2-cell stages. In Yan dataset (2), we did not find such pattern and cells from oocyte to blastocyst stages were included in the analysis. SCnorm failed to converge in the number of gene groups in Deng dataset and was not reported. Goolam dataset (3) contains 4 batches with different dilution of ERCC spike-ins. We analyzed all cells when estimating effective library size by different normalization methods but only included cells from batch 1 when estimating absolute mRNA content by ERCC spike-ins.

Differential expression analysis was carried out using R package DESeq2 (4) by feeding DESeq2 with size factors estimated from different methods. Comparison was conducted between late 2-cell *vs.* middle 2-cell, 4-cell *vs.* late 2-cell, 8-cell *vs.* 4-cell and 16-cell *vs.* 8-cell stages. Genes with false discovery rate less than 0.05 or 0.15 were considered as DE genes.

We combined scRNA-seq data (1) and ATAC-seq data (5) to define a set of maternal genes. Firstly genes with average ${log}_{2}(TPM+1)$ value large than 3 in cells from zygote stage and less than 1 in cells from 16-cell and blastocyst stages in Deng dataset were defined as maternal genes. We tried to remove false maternal genes based on their ATAC-seq signal in promoter regions (measured by average rpm bp^-1^ value across the promoter region). Promoter regions were defined as upstream and downstream region within 2kb of transcription starting site (TSS). We picked a set of transcriptional quiescent genes with zero TPM value across all cells in Deng dataset as negative control. Considering that there might be some outliers in quiescent genes (with open chromatin in promoter region but not detected by scRNA-seq), we defined a threshold as 90% quantile of ATAC-seq signal of all quiescent genes. We removed maternal genes if their ATAC-seq signal exceeded this threshold in 2-cell or 4-cell stages. Regardless of the normalization methods applied, most maternal genes were down-regulated, suggesting that our maternal gene list was reliable. We checked the fold change of maternal genes between late 2-cell vs. middle 2-cell and 4-cell vs. late 2-cell stages as we thought their expression were relatively high in these stages and less affected by our definition (we constrained the average ${log}_{2}(TPM+1)$ value of maternal genes in 16-cell stage below 1).

#### 1.2 Mouse embryogenesis ATAC-seq data processing

Raw reads from mouse embryogenesis ATAC-seq data (5) were first processed by Trim Galore to remove adapters and then aligned to *M. musculus* genome (Ensembl v.38.89) using Bowtie2 v2.3.4.1 with the following parameters: (--t --q --N1 --L 25 --X 2000 --no-mixed --no-discordant). Bam files of the same developmental stage were merged by Samtools-1.8 (6). Reads coverage was counted by IGVtools v2.3.98 (Broad Institute) (7) with the following parameters: (-w 100 -e 250) and normalized to rpm bp^-1^. The Galaxy deeptools suite (8) was used to plot heat maps.

#### 1.3 Patel dataset processing

RNA sequencing data processing and Copy number inference was conducted on TPM matrix of Patel dataset (9) following the instructions of original article. PCA was carried out using R function prcomp with no scaling on standard deviation on log2 transformed normalized matrix by different methods after adding one pseudo count. The dominance of CDR on PC1 and PC2 was measured by coefficient of determination (*R*^2^) from a linear regression model (CDR *~* PC1 + PC2) using R function lm. The effective library size of malignant cells was calculated as described above. The relationship between PC1 and effective library size of malignant cells upon ISnorm was measured by linear regression between log transformed effective library size and PC1 (log effective library size ~ PC1) using R function lm as we found the linear relationship can be better modeled after transformation (data not shown). Genes with top 100 absolute loadings in PC1 and PC2 respectively were referred to as high loading genes. Oligodendrocyte markers and up-regulated genes in radial glia cells were obtained from MSigDB v6.2 (10). The difference of CDR and effective library size between two tumor clones and normal cells were tested by Analysis of Variance (ANOVA) using R function aov. Four cells with significantly high effective library size (larger than 500) were considered as low quality cells and were excluded in boxplot and ANOVA.

### 2 Supplementary Figures

#### 2.1 Supplementary Figure S1

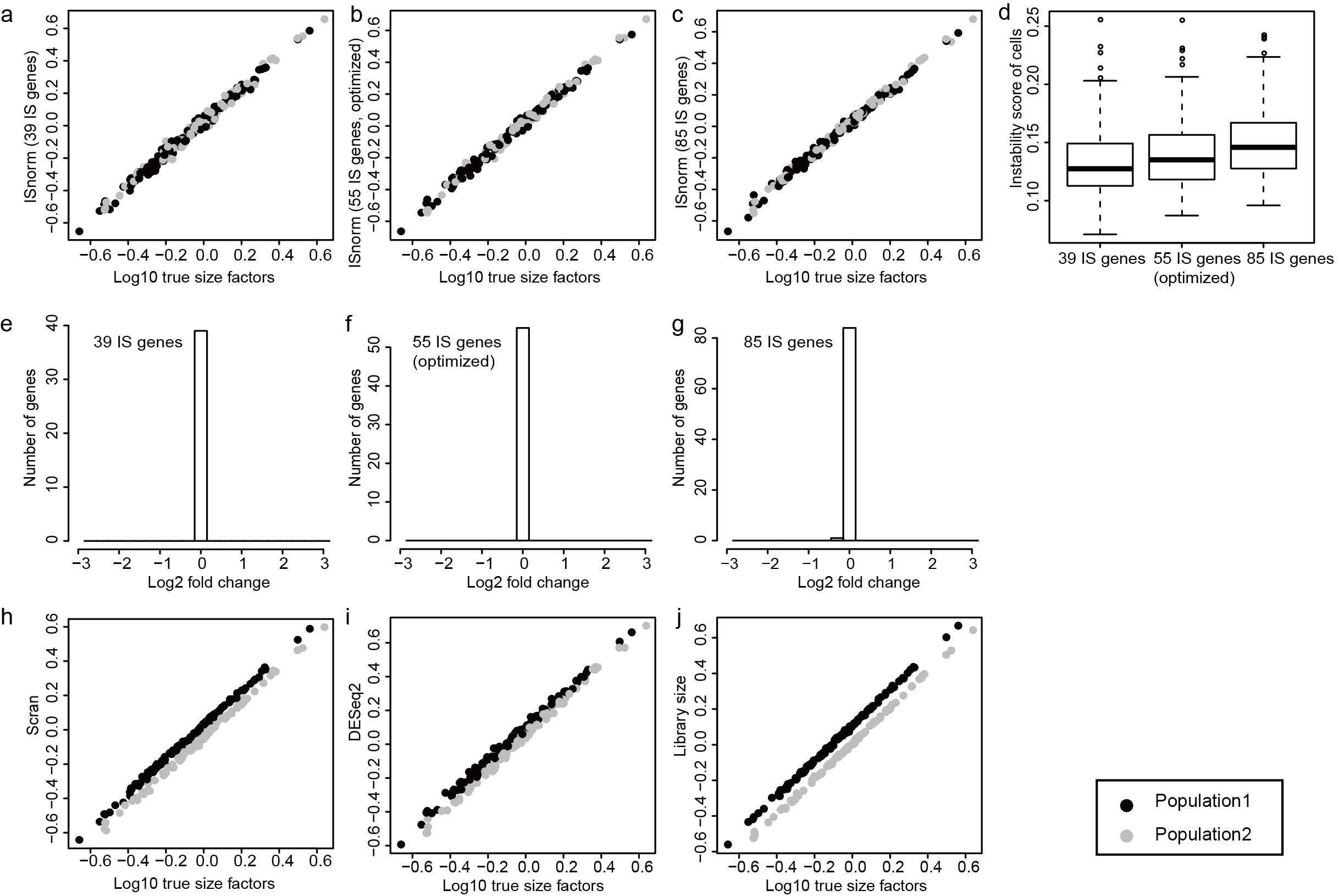

**Figure S1 Performance of ISnorm and other existing methods on inDrop simulation.** a-c Size factors estimated by ISnorm. **d** Instability scores of cells for each candidate set. **e**-**g** Distribution of log2 fold change of IS genes in each candidate set. **h**-**j** Size factors estimated by other existing methods. SCnorm failed to calculate a normalized matrix in inDrop simulation and was not reported.

#### 2.2 Supplementary Figure S2

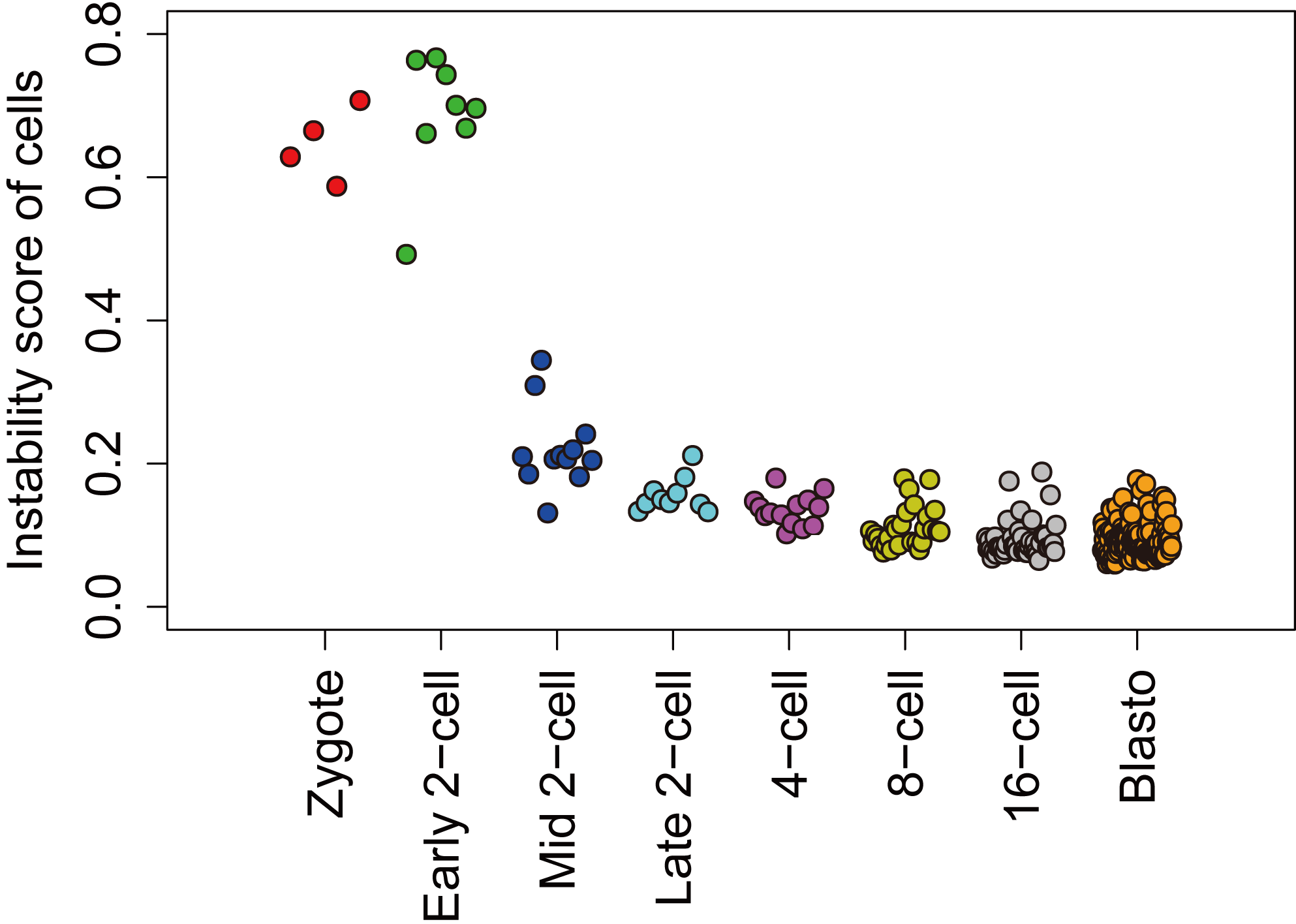

**Figure S2 Instability scores for cells in Deng dataset of embryogenesis, showing cells in zygote and early 2-cell stages had higher instability scores.**

#### 2.3 Supplementary Figure S3

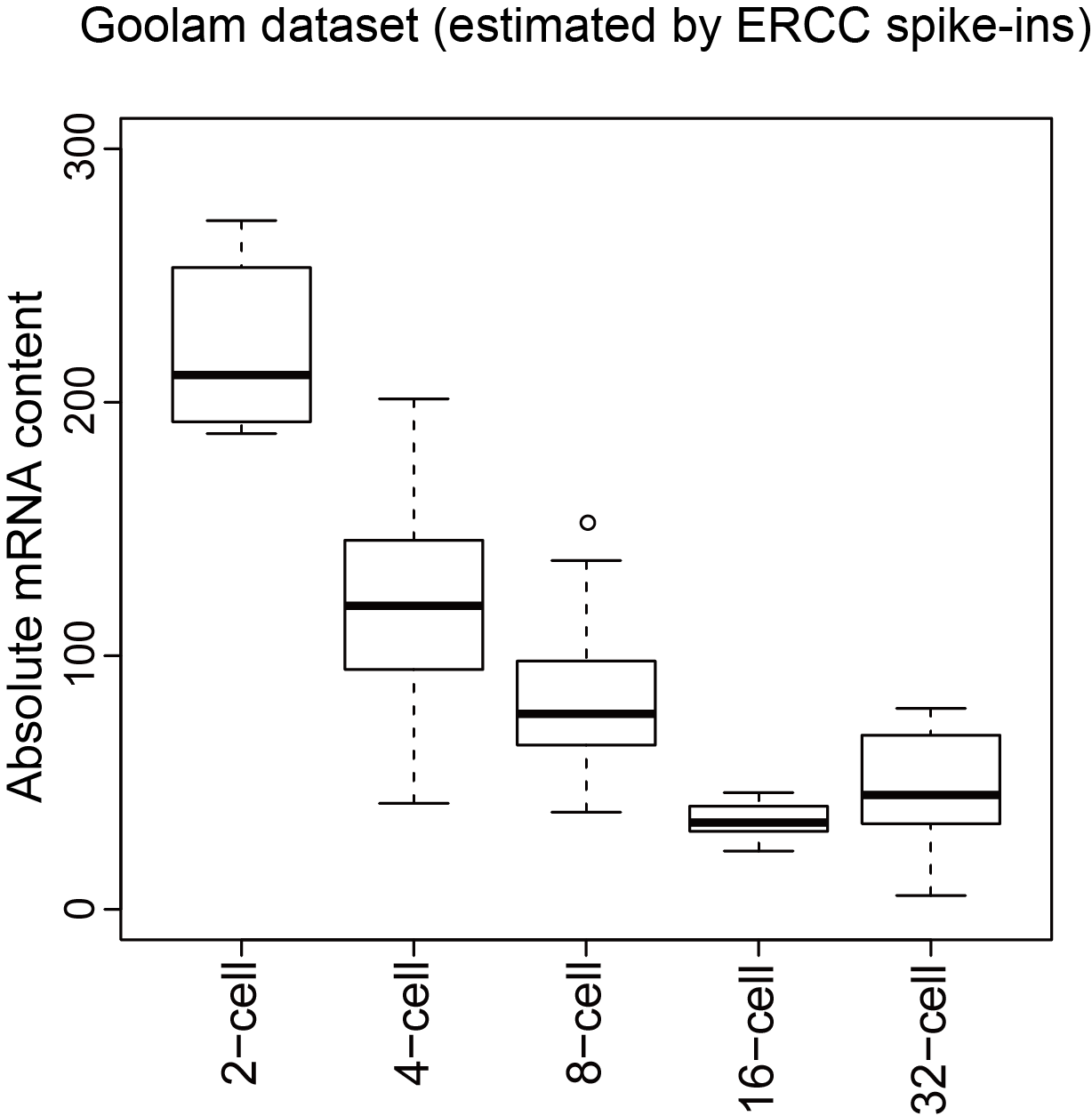

**Figure S3 Absolute mRNA content estimated by ERCC spike-ins on Goolam dataset for all cells from batch 1.**

#### 2.4 Supplementary Figure S4

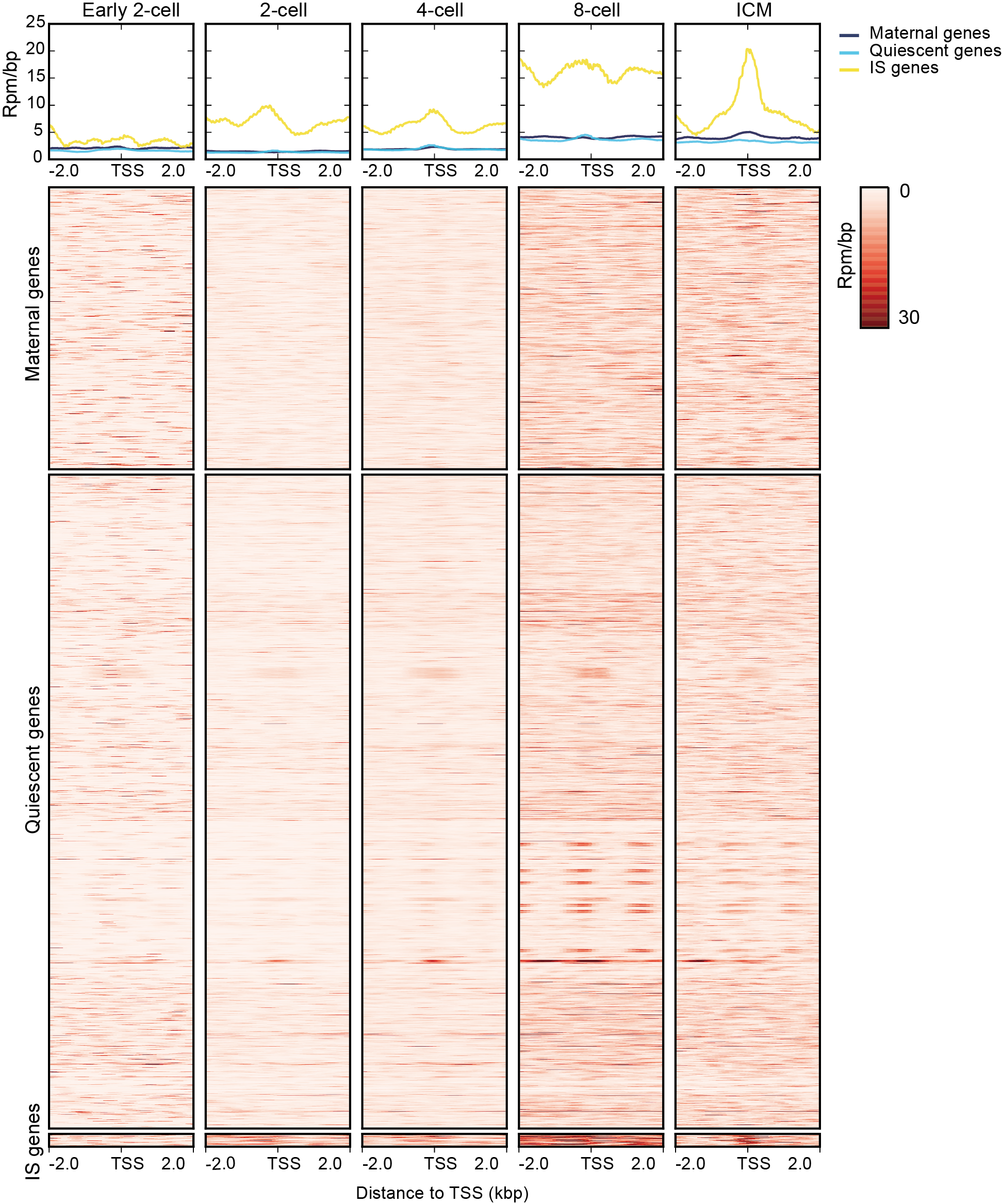

**Figure S4 ATAC-seq signal in promoter region of selected maternal genes, quiescent genes and IS genes from ATAC-seq data of mouse embryogenesis.**

#### 2.5 Supplementary Figure S5

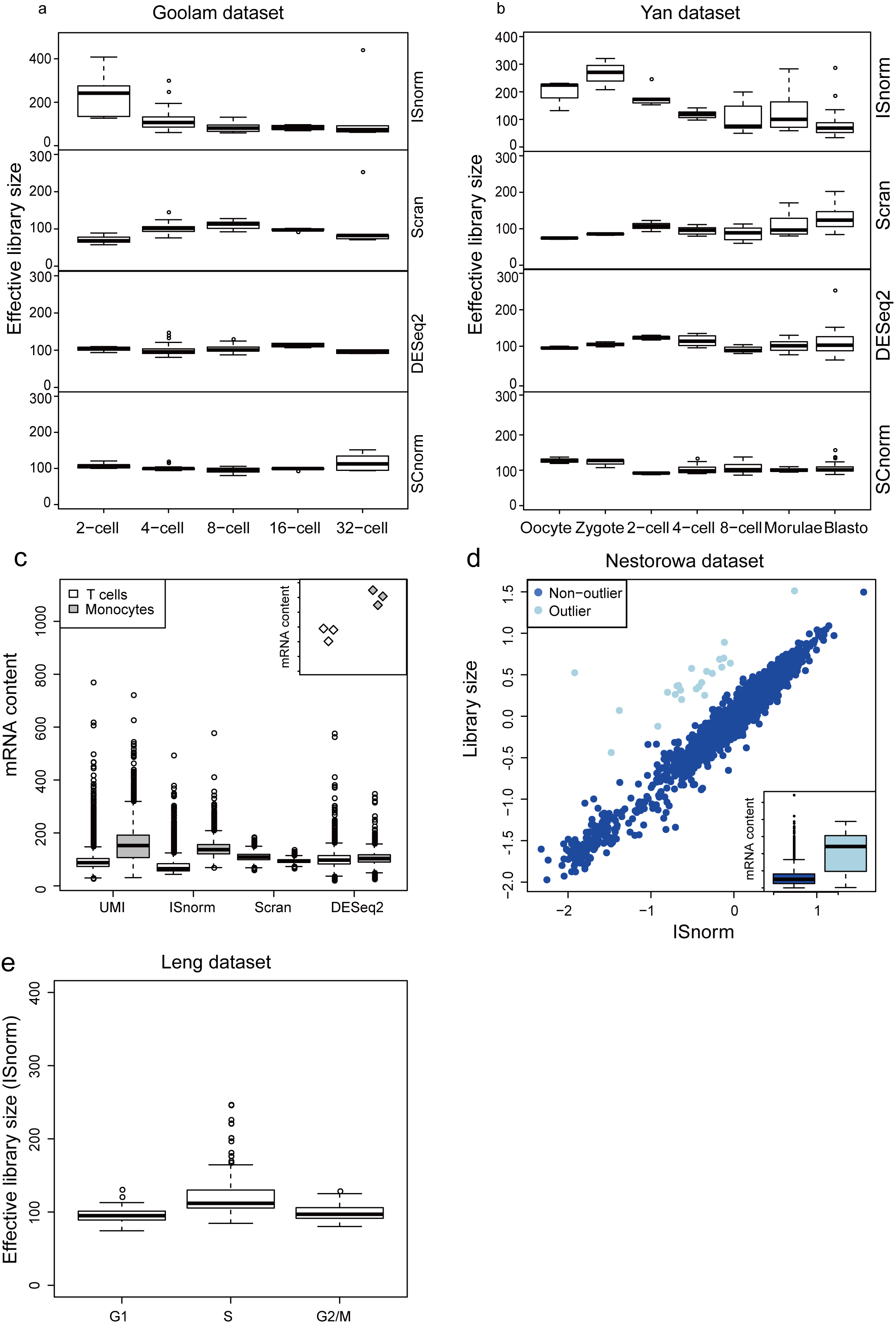

**Figure S5 ISnorm estimates mRNA content difference among single cells.** **a** Goolam dataset. **b** Yan dataset. **c** Comparison of total UMI counts and effective library size by different normalization methods on PBMC datasets. Qualification of mRNA content for T cells and monocytes by ERCC spike-ins through bulk RNA-seq are shown in upper right. **d** Comparison of size factors by ISnorm and library size on Nestorowa datasets, showing outlier cells with higher mRNA content. Absolute mRNA content estimated by ERCC spike-ins for outliers and non-outliers are shown in lower right. **e** Effective library size for cells in G1, S and G2/M stages estimated by ISnorm on Leng dataset.

#### 2.6 Supplementary Figure S6

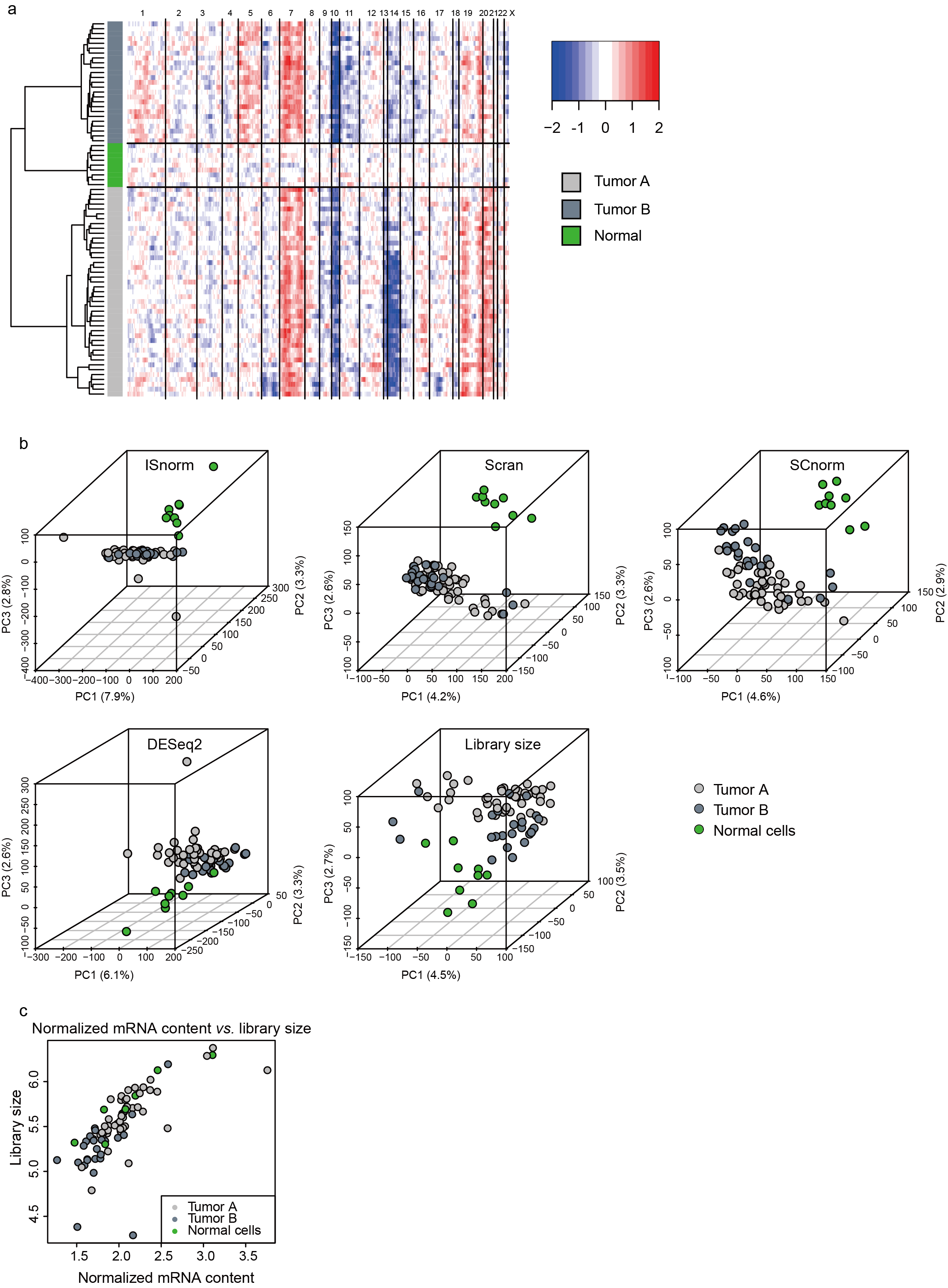

**Figure S6 Benchmarking on scRNA-seq of glioblastoma of patient MGH31 from Patel dataset.** **a** Copy number variation analysis inferring normal cells and tumor clones. **b** The results of first three PCs inferred by PCA on normalized matrices upon ISnorm and other existing methods. **c** Comparison between library size and normalized mRNA content upon ISnorm.

#### 2.7 Supplementary Figure S7

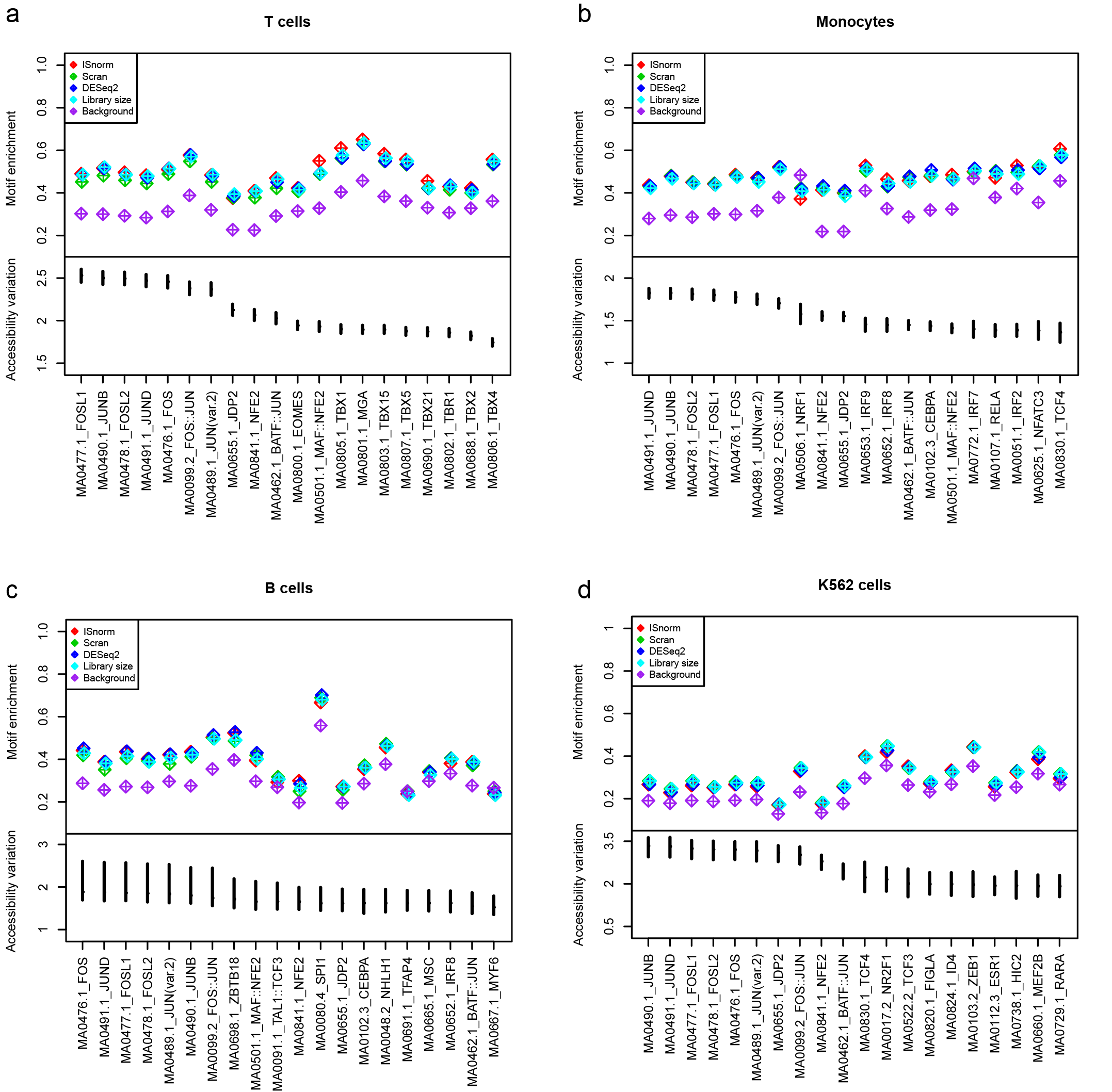

**Figure S7 The results of motif enrichment analysis.** **a** T cells, **b** monocytes, **c** B cells and **d** K562 cells. The upper part shows the pattern of motif enrichments based on ISnorm and other normalization methods and the lower part shows the point estimates and error bars of accessibility variation of motifs from scATAC data.

#### 2.8 Supplementary Figure S8

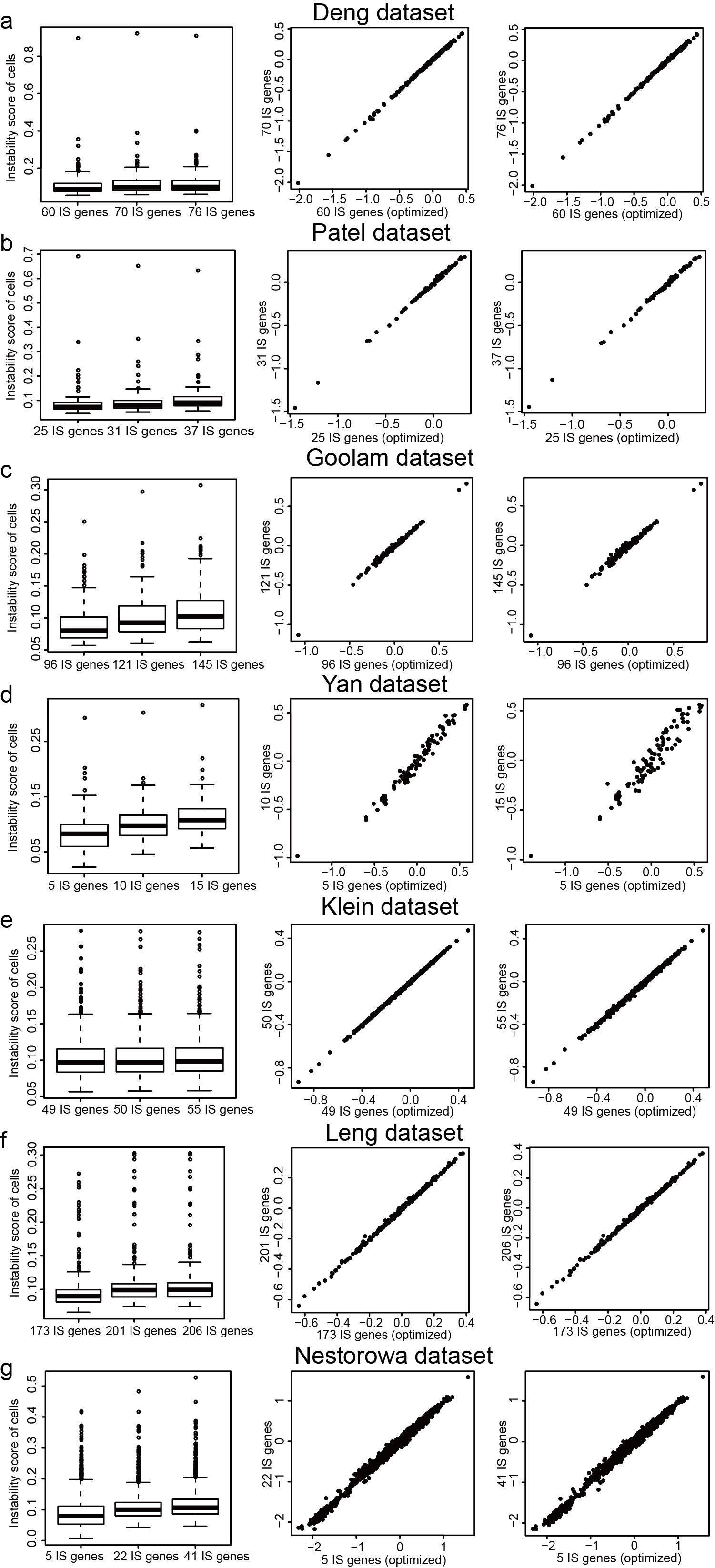

**Figure S8 Comparison of instability scores and size factors estimated by different candidate sets**. **a** Deng dataset, **b** Patel dataset, **c** Goolam dataset, **d** Yan dataset, **e** Klein dataset, **f** Leng data set and **g** Nestorowa dataset. Figures in the first column show the comparison between instability scores and figures in the second and third columns show the comparison between size factors.

#### 2.9 Supplementary Figure S9

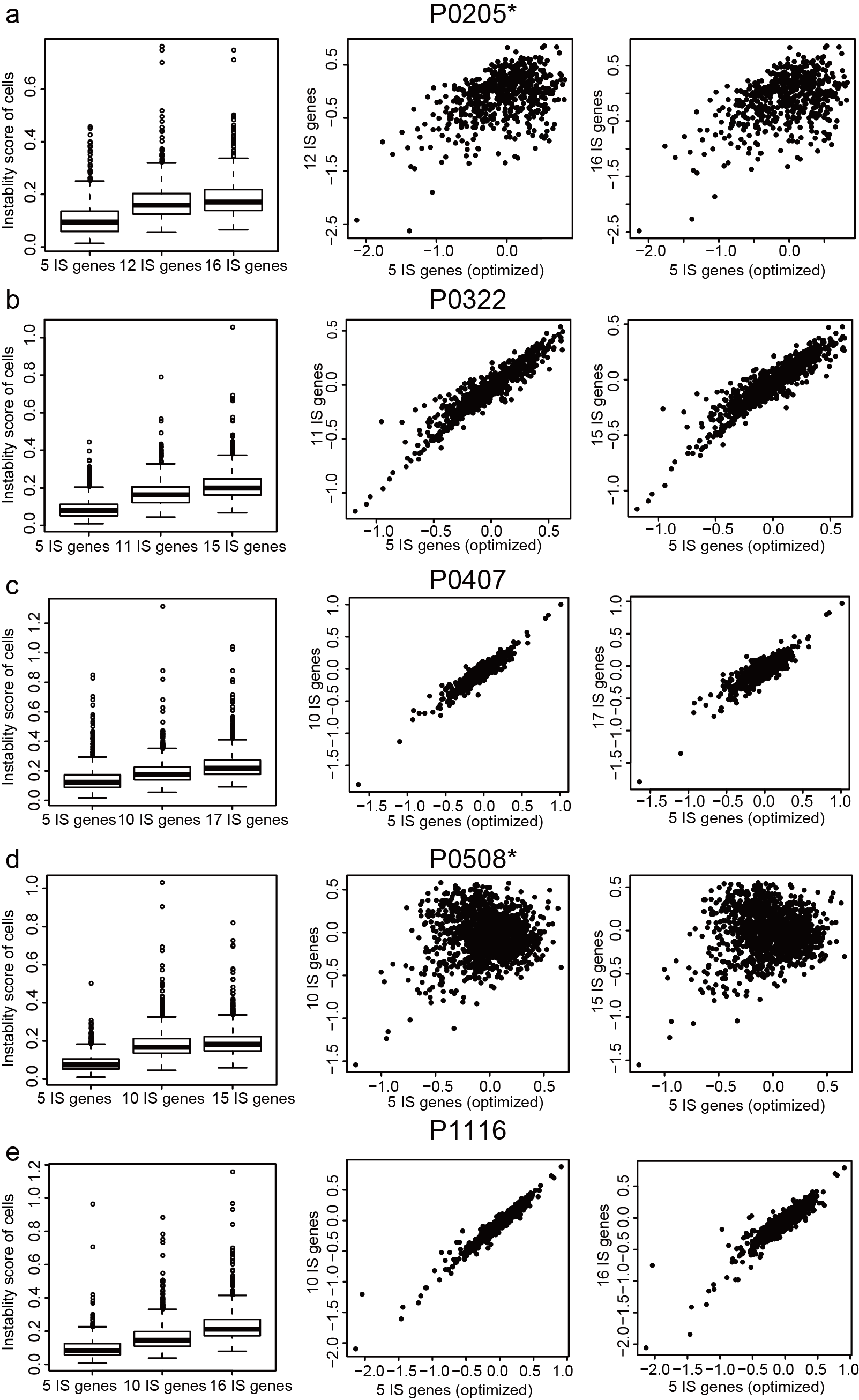

**Figure S9 Comparison of instability scores and size factors estimated by different candidate sets for Zheng dataset of samples.** **a** P0205, **b** P0322, **c** P0407, **d** P0508 and **e** P1116. Figures in the first column show the comparison between instability scores and figures in the second and third columns show the comparison between size factors. Sample P0205 and P0508 are marked by an asterisk, showing that optimized geneset does not share common genes with other candidate genesets.

#### 2.10 Supplementary Figure S10

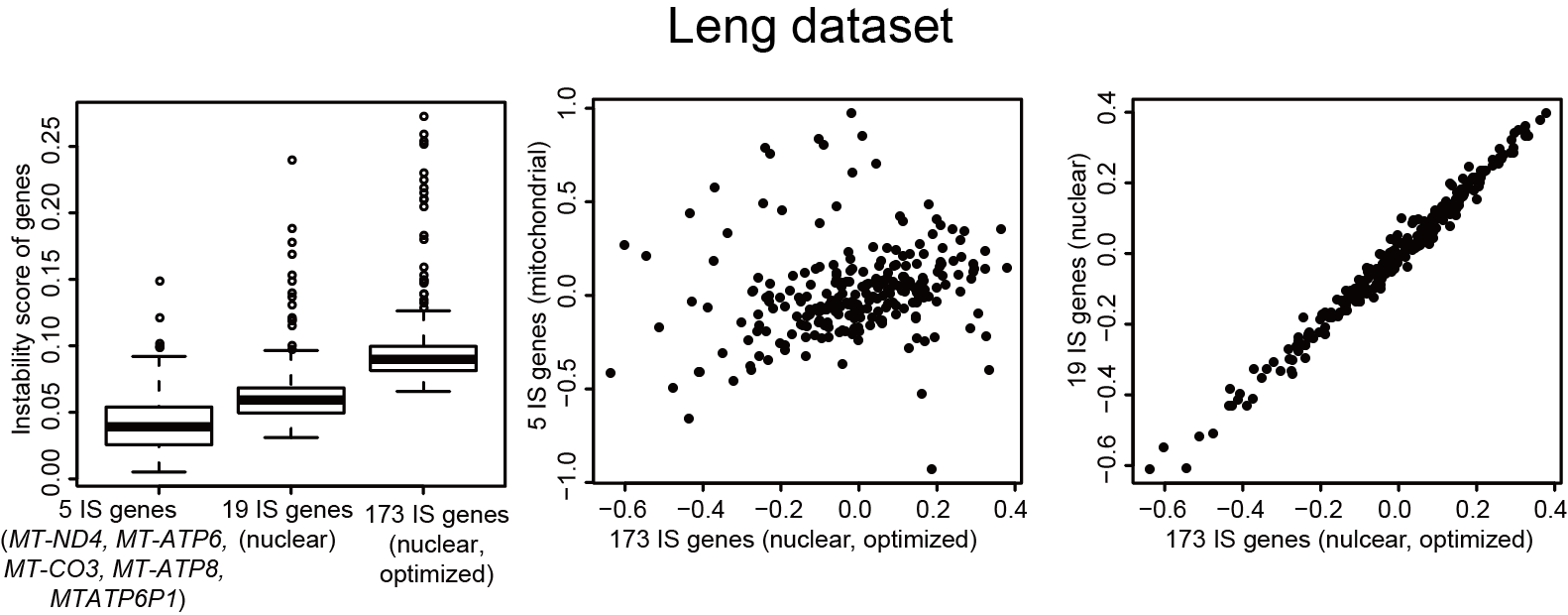

**Figure S10 Comparison of instability scores and size factors estimated by mitochondrial geneset and nuclear genesets for Leng dataset.** Figures in the first column show the comparison between instability scores and figures in the second and third columns show the comparison between size factors. The results of mitochondrial geneset, one non-optimal nuclear geneset with 19 genes and the optimal geneset with 173 genes are shown.

#### 2.11 Supplementary Figure S11

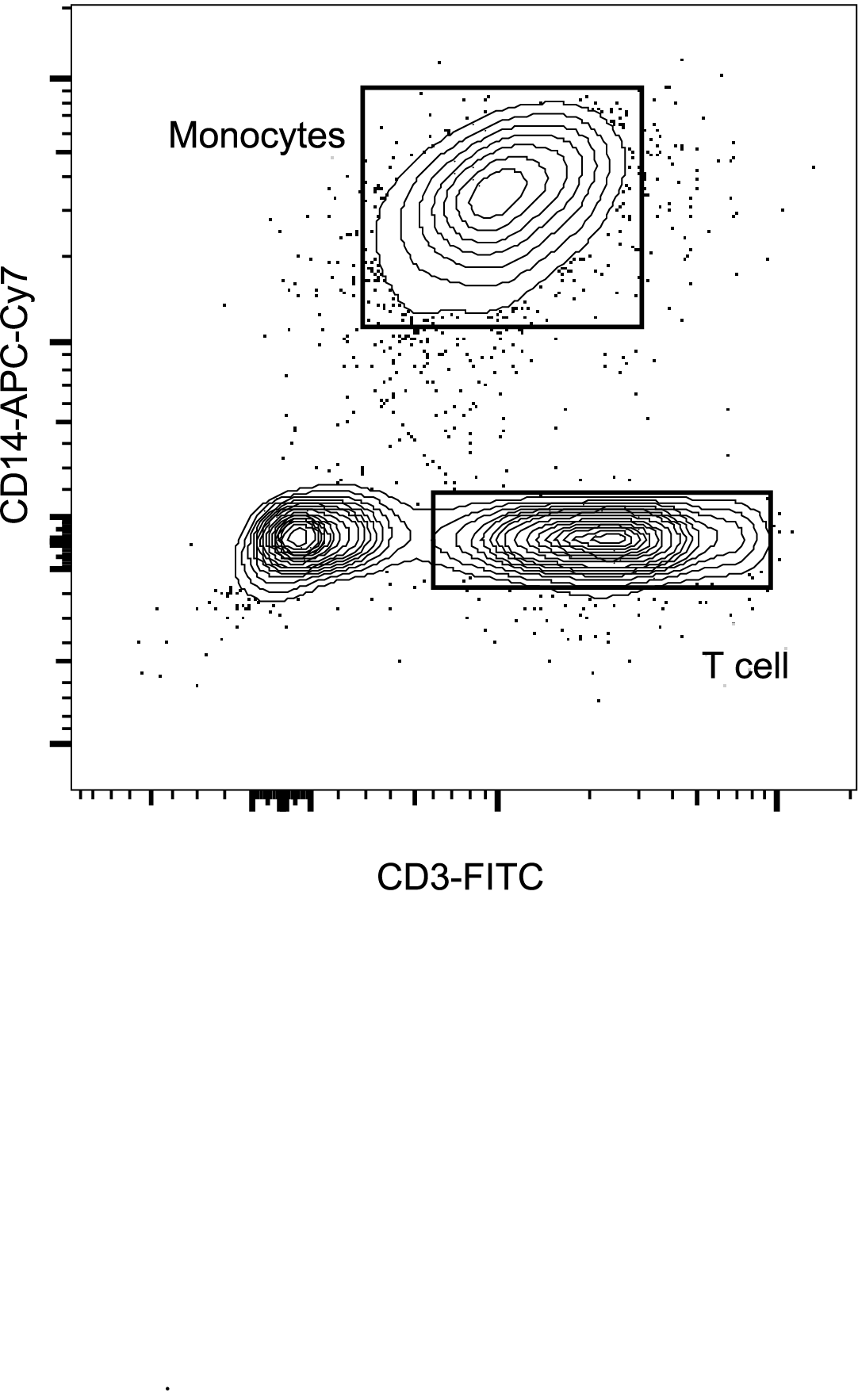

**Figure S11 FACS analysis of monocytes and T cells.** Isolated PBMC was stained with human CD3 and CD14 antibody, sorting for the same number of monocytes and T cells as shown.

#### 2.12 Supplementary Figure S12

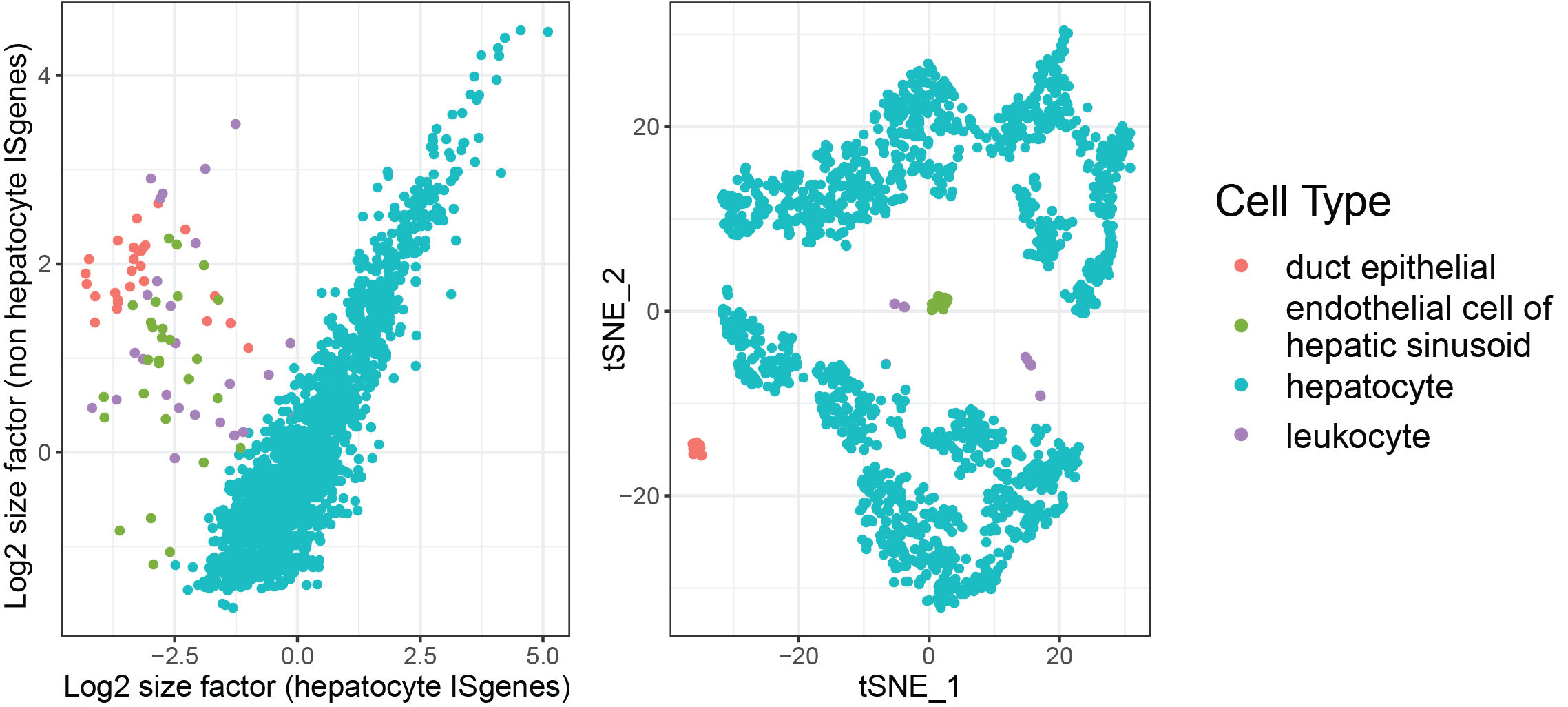

**Figure S12 ISnorm fails when cell populations do not share common IS genes.** Figure in the left shows the comparison of size factors learned from hepatocytes and non hepatocytes (see Liver Hepatocyte and Liver Non-Hepatocyte from 10X protocol in Table S3). IS genes learned from different cell types do not agree with each other. Figure in the right shows the corresponding cell types in liver tissue.

### 3 Supplementary Tables

#### 3.1 Supplementary table S1

**Table S1 datasets used in this study.**

| Dataset | Accession number | Protocol | Raw data | Biotype |
| --- | --- | --- | --- | --- |
| Deng et al. (1) | GSE45719 | Smart-seq | raw reads | Mouse embryogenesis |
| Patel et al. (9) | GSE57872 | Smart-seq | raw reads | Glioblastoma |
| Goolam et al. (3) | E-MTAB-3321 | Smart-seq2 | raw reads | Mouse embryogenesis |
| Yan et al. (2) | GSE36552 | Tang et al. (11) | raw reads | Human embryogenesis |
| Klein et al. (12) | GSE65525 | inDrop | processed raw counts | Mouse ESCs |
| Klein et al. (12) | GSE65525 | inDrop | processed raw counts | K562 cells |
| Leng et al. (13) | GSE64016 | Smart-seq | raw reads | Human ESCs |
| Nestorowa et al. (14) | GSE81682 | Smart-seq2 | processed raw counts | Mouse HSPCs |
| Zheng et al. (15) | GSE98638 | Smart-seq | processed raw counts | T cells (P0205, P0322, P0508, P1116) |
| Zheng et al. (15) | GSE98638 | Smart-seq | processed raw counts | T cells (P0407) |
| Bach et al. (16) | GSE106273 | 10X Genomics Chromium | processed raw counts | Mammary epithelial cells |
| PBMCs |  | 10X Genomics Chromium  (scRNA-seq) | processed raw counts | Human PBMCs from a healthy donor |
| Tabula Muris. (17) | GSE109774 | 10X Genomics Chromium and  Samrt-seq2 | processed raw counts | Mouse organs and tissues |
| PBMCs |  | 10X Genomics Chromium  (scATAC-seq) | - | Human PBMCs from a healthy donor |
| Wu et al. (5) | GSE66390 | ATAC-seq | - | Mouse embryogenesis |
| Ziegenhain et al. (18) | GSE75790 | Smart-seq2 | - | Mouse ESCs |
| Schep et al. (19) | GSE99172 | Single-cell ATAC-seq | - | K562 cells |
| This paper | HRA000048 | Bulk RNA-seq |  | Human CD3+ T cells and CD14+ monocytes |

#### 3.2 Supplementary table S2

**Table S2 IS genes selected for benchmarking dataset.**

Genes that are known to be up-regulated in radial glia cells from MSigDB in Patel dataset are

indicated by the asterisk.

| Dataset | IS genes |
| --- | --- |
| Deng et al. (1) | *Rps11, Rpl8, Rps9, Rps18, Rps15a, Rps25, Rpl15, Rps5, Rpl19, Rps27a, Rps14, Rps24, Rps26, Rplp2, Rps3a1, Rps20, Rps6, Rpl6, Rps3, Rpl4, Rps16, Rps21, Rps19, Rpl37, Rpl7, Rpl18a, Rpl37a, Rpl9, Rps8, Rpl29, Rpl32, Gm10036, Rpl23a, Rpl18, Rpl11, Rps26-ps1, Rps16-ps2, Dppa5a, Rpl26, Naca, Rps7, Rps17, Rps3a2, Rpl35, Rps15, Rps28, Rpl23, Ppia, Rpl37rt, Rpl13a, Gm2000, Gm8172, Rps27, Gm6472, Rpl24, Rps23-ps1, Gm5835, Gm4332, Rpl7-ps7, Gm9794* |
| Patel et al. (9) | *BLOC1S6, FRK*, MORC4*, TBC1D8B*, KIAA1328*, ADAMTSL3, ATP9B, CYSLTR1, SULT1B1*, C4orf32*, TMEM45A, TMEM212*, C12orf55*, AL590867.1, XKR9*, XXbac-BPG154L12.4, RP11-47I22.3, AC007743.1, LINC00504, AL139147.1, UGDH-AS1, RP11-703G6.1, RP11-259O2.3, RP11-946L20.4, RP11-346D14.1* |
| Goolam et al. (3) | *Rpl13, Rps11, Rpl8, Psmd4, Psmb4, Rps9, Rplp1, Rps15a, Rps25, Rpl15, Rps5, Psmb1, Rpl19, C1qbp, Rack1, Rps27a, Psmc5, Nme2, Psma6, Btf3, Hsp90ab1, Rps14, Rps24, Rps26, Atp5b, Rplp2, Psmb7, Tpm3, Rps20, Rps6, Rpl22, Tomm7, Rpl6, Pomp, Cct7, Rpl28, Rps3, Cox7a2, Rpl4, Rps29, Eef2, Cox6b1, Rps16, Rpl10a, Fau, Rps21, Adrm1, Rps19, Rpl37, Rpl7, Rpl18a, Rpl37a, Rpl27a, Rpl9, Rps8, Rpl29, Rpl36al, Rps10, Eef1d, Cfl1, Gm7536, Snrpg, Rpl38, Rpl32, Rpl36, Rpl23a, Rpl30, Rpl28-ps1, Rpl18, Rpl11, Rpl35a, Naca, Rps17, Rpl34, Rpl17, Rpl7a, Rpl35, Rpl27, Rps15, Rps28, Rpl31-ps8, Gm10275, Rpl23, Ppia, Rpl37rt, Rpl27-ps3, Rpl31, Rpl13a, Gm2000, Eif5a, Gm15500, Uba52, Rps27, Rps13, Rpl41, Rpl24* |
| Yan et al. (2) | *RPS16, RPS25, RPL32, RPL27A, RPL35A* |
| Klein et al. (12) | *Eef1a1, Gm10653, Gm5779, Gm6548, Rpl12, Rpl13, Rpl13a, Rpl14, Rpl14-ps1, Rpl15, Rpl18a, Rpl23, Rpl27a, Rpl31-ps12, Rpl38, Rpl41, Rpl8, Rplp0, Rplp1, Rplp2, Rplp2-ps1, Rps12, Rps16, Rps18, Rps19, Rps2, Rps24, Rps3a1, Rps4x, Rps5, Rps8, Rpsa, Tpt1* |
| Klein et al. (12) | *EEF1A1, EEF1G, KLF15, MIR1234, RPL10, RPL13, RPL14, RPL15, RPL18, RPL22, RPL23, RPL27A, RPL3, RPL35, RPL4, RPL41, RPL5, RPL6, RPL7, RPL7A, RPLP0, RPLP1, RPS12, RPS15, RPS18, RPS19, RPS2, RPS23, RPS28, RPS5, RPS7* |
| Leng et al. (13) | *PSMB1, MGST1, RPS20, TMSB10, PSMA4, THRAP3, RPL18, RPL31, ENO1, ACTB, XRCC5, HSP90AA1, RPS5, GSTP1, FTL, RPL6, RPLP0, HNRNPC, MYL6, HSP90AB1, RPL3, EIF3E, RPS16, RPS19, RPL18A, SSBP1, RPL19, YWHAE, RPL34, RPS13, GAPDH, TPI1, LDHB, SRSF3, RPS12, GMNN, COX7A2, SKP1, RPL24, EEF1B2, NCL, RPS15, HSPE1, PRDX1, STMN1, RPS25, SET, SLIRP, RPL21, RPL5, HNRNPA2B1, TUBA1B, RPS10, RPL23, SNRPD2, COX7C, SHFM1, PAICS, RPL36, COX4I1, RPL27, RAN, H3F3B, POMP, TPT1, ATP5G2, HNRNPA1, TXN, RPL35, RPS6, HMGA1, RPLP1, RPS24, SSB, SNRPF, SERF2, SRP14, RPS11, RPL13A, RPL11, RPS8, CALM2, RPS27A, SNRPG, HSPD1, RPL32, RPS3A, RPL37, RPL10, RPL7, RPL7A, RPS3, FAU, ALDOA, DBI, RPL30, EEF1A1, RPL8, LSM12, RPL26, RPL29, H3F3A, RPL9, H2AFZ, RPS14, FABP5, COX6C, YWHAZ, RPL36AL, PRDX3, RPL27A, NGFRAP1, NDUFS5, ATP5I, HINT1, UBB, RSL1D1, RPS21, RPS7, RPL38, UQCRH, RPL4, RPL15, RPLP2, RPS27, NPM1, NOP10, RPL35A, CD24P4, MORF4L1, RPS23, NAP1L1, PTMA, H3F3C, SUMO2, RPL14, NDUFA4, HMGB1, TUBB, PPIA, NACA, TOMM7, RPL37A, RPL12, RPL23A, RPL10A, HMGN2, RPL39, RP11-217O12.1, GNB2L1, TMSB4X, RP4-765C7.2, RPL12P38, RPS29, PTMAP5, UBA52, HNRNPA1P48, RP11-742N3.1, OST4, RPL41, CTB-63M22.1, RPL10P3, RPS18, TMA7, RPS28, H3F3AP4, L1TD1, ATP5J2, ATP5O, EEF1G, RP11-386G11.10, RPL17, RP5-940J5.9* |
| Nestorowa et al. (14) | *Rps18-204, Rps19, Rpl32, Rps19-ps6, Gm7266* |
| Zheng et al. (15) | *HLA-A, HLA-B, HLA-G, HLA-H, HLA-J* |
| Zheng et al. (15) (P0407) | *HLA-B, HLA-C, HLA-G, HLA-L, B2M* |
| Bach et al. (16) | *Rpl10, Rps4x, Rps3a1, Rps8, Rpl9, Rpl32, Rps5, Rps19, Rps16, Rps11, Rpl13a, Rps3, Rps15a, Rplp2, Rpl18a, Rpl13, Rps27a, Rpl26, Rpl23, Rpl19, Rps23, Rps7, Rpl37, Rpl8, Rps2, Rps18, Rps14, Rpl17* |
| PBMCs | *RPL11, RPS8, RPS27A, RPL32, RPL29, RPL35A, RPL9, RPL34, RPS3A, RPL37, RPS23, RPS14, RPS18, EEF1A1, RPS12, RPS4X, RPL39, RPL10, RPL30, RPS6, RPL7A, RPLP2, RPL41, RPL21, TPT1, RPS2, RPS15A, RPL13, RPL26, RPL19, RPS15, RPS28, RPL18A, RPL18, RPL28* |

#### 3.3 Supplementary table S3

**Table S3 IS genes selected among different tissues.**

| Tissue | Protocol | IS genes |
| --- | --- | --- |
| Aorta | Smart-seq2 | *Rn45s,Rpl13,Rpl41,Rps29,Tpt1* |
| Bladder | Smart-seq2 | *Rpl13,Rpl18a,Rplp0,Rps11,Rps14,Rps18,Rps19,Rps9* |
| Brain_Myeloid | Smart-seq2 | *C1qa,C1qb,C1qc,Cst3,Ctsd,Ctss,Hexb,Itm2b,Lgmn,Selplg,Tmsb4x* |
| Brain_Non_Myeloid | Smart-seq2 | *Cfl1,Gm1821,Oaz1,Ppia,Ubb* |
| Diaphragm | Smart-seq2 | *Mir682,Rpl13,Rpl18a,Rpl36,Rpl37,Rpl37a,Rpl41,Rps14,Rps16,Rps18,*  *Rps19,Rps28,Rps29* |
| Fat | Smart-seq2 | *Mir682,Rpl18a,Rpl35,Rpl36,Rpl37,Rpl37a,Rpl41,Rplp2,Rps14,Rps19,*  *Rps28,Rps29* |
| Heart_Aorta | Smart-seq2 | *Rpl35,Rpl37,Rpl37a,Rpl41,Rps19,Rps27,Rps28,Rps29* |
| Heart | Smart-seq2 | *Rpl35,Rpl37,Rpl37a,Rpl41,Rps19,Rps28,Rps29* |
| Kidney | Smart-seq2 | *Gm1821,Ppia,Rn45s,Rpl41,Ubb* |
| Large_Intestine | Smart-seq2 | *Mir682,Rpl28,Rpl36,Rpl37,Rpl37a,Rpl38,Rpl41,Rps28,Rps29,Uba52* |
| Limb_Muscle | Smart-seq2 | *Eef1a1,Mir682,Rpl13,Rpl23a,Rpl32,Rpl35,Rpl36,Rpl37,Rpl37a,Rpl41,*  *Rplp1,Rplp2,Rps14,Rps19,Rps23,Rps25,Rps27,Rps28,Rps29* |
| Liver_Hepatocyte | Smart-seq2 | *Apoa2,Apoc1,Atp5l,Cox6c,Ddt* |
| Liver_Non_  Hepatocyte | Smart-seq2 | *B2m,Gm1821,H2-D1,Lars2,Rn45s* |
| Lung | Smart-seq2 | *Eif1,Gm1821,Oaz1,Ppia,Ubb* |
| Mammary_Gland | Smart-seq2 | *Rpl13,Rpl18a,Rpl32,Rpl37,Rps14,Rps18,Rps19,Rps23* |
| Marrow | Smart-seq2 | *Mir682,Rpl34-ps1,Rpl37,Rpl37a,Rpl41,Rps19,Rps28,Rps29,Rps9* |
| Pancreas | Smart-seq2 | *Mir682,Rpl36,Rpl37,Rpl37a,Rpl38,Rpl41,Rps14,Rps21,Rps28,Rps29* |
| Skin | Smart-seq2 | *Rpl13,Rpl18a,Rps14,Rps18,Rps19* |
| Spleen | Smart-seq2 | *Mir682,Rpl13,Rpl13a,Rpl18a,Rpl36,Rpl37a,Rpl41,Rps19,Rps27,Rps28,*  *Rps29* |
| Thymus | Smart-seq2 | *Mir682,Rpl37a,Rpl41,Rps14,Rps27,Rps29* |
| Tongue | Smart-seq2 | *Rpl13,Rpl15,Rpl18a,Rpl23a,Rpl32,Rpl35,Rpl36,Rpl37a,Rpl41,Rpl9,Rplp0,Rplp1,Rplp2,Rps11,Rps14,Rps16,Rps18,Rps19,Rps20,Rps3,Rps3a,Rps7,Rps9,Rpsa* |
| Trachea | Smart-seq2 | *Mir682,Rpl35,Rpl36,Rpl37,Rpl37a,Rpl41,Rplp1,Rps23,Rps28,Rps29* |
| Bladder | 10X | *Rpl37a,Rps15,Rpl41,Rps27a,Rpl23a,Rpl23,Rpl19,Rps7,Rps29,Rps23,Rps24,Rpl37,Rpl8,Rpl3,Rpl35a,Rpl24,Rps2,Rpl10a,Rps28,Rps18,Rpl36,*  *Rps14,Rpl17,Fau,Rpl35,Rps3a,Rps6,Rps8,Rpl11,Rpl9,Rplp0,Rpl6,Rpl21,Rpl32,Rps9,Rps5,Rps19,Rps16,Rps11,Rpl13a,Rps3,Rpl27a,Rps13,*  *Rps15a,Rplp2,Uba52,Rpl18a,Rpl13,Rps25,Rplp1,Eef1a1,Rpl14,Rpl39,*  *Rpl10,Rps4x* |
| Heart_Aorta | 10X | *Rpl37a,Rpl41,Rpl23a,Rps29,Rps23,Rps24,Rpl37,Rpl8,Rpl35a,Rps14,*  *Rpl35,Rpl9,Rpl21,Rpl32,Rps5,Rps19,Rps16,Rpl13a,Rplp2,Uba52,Rpl18a,Rpl13,Rplp1,Rpl14,Rps4x* |
| Kidney | 10X | *Rpl37a,Rpl41,Rpl23,Rpl38,Rps29,Rpl37,Rpl35a,Rpl36,Rpl35,Rpl32,*  *Rplp2,Uba52,Rplp1,Eef1a1,Rpl39* |
| Limb_Muscle | 10X | *Rpl7,Rpl37a,Rps12,Rps15,Rpl41,Rps27a,Rpl23a,Rpl23,Rpl19,Rpl38,Rps7,Rps29,Rps23,Rps24,Tpt1,Rpl37,Rpl8,Rpl35a,Rps10,Rps28,Rps18,Rpl36,Rps14,Rpl17,Fau,Rpl35,Rps3a,Rps20,Rps6,Rps8,Rps15a-ps4,Rpl11,Rpl9,Rplp0,Rpl6,Rpl21,Rpl32,Rps9,Rps5,Rps19,Rps16,Rps11,Rpl13a,Rpl18,*  *Rps17,Rps3,Rpl27a,Rps13,Rps15a,Rplp2,Uba52,Rpl18a,Rpl13,Rps25,*  *Rplp1,Eef1a1,Rpl14,Rpl39,Rps4x* |
| Liver_Hepatocyte | 10X | *Apoa2,Serpina1b,Serpina1a,Serpina1c,Ttr,Apoc4,Apoc1,Ftl1,Apoa1,*  *Apoc3* |
| Liver_Non  Hepatocyte | 10X | *Rpl37a,Rps15,Rpl41,Gnb2l1,Rpl37,Rpl3,Rpl35a,Rps14,Rpl17,Fau,Rpl35,Rps20,Rpl32,Rpl28,Rps5,Rps19,Rps16,Rps11,Rpl13a,Rpl18,Rpl27a,*  *Rpl18a,Rps25,Rplp1,Rps4x* |
| Lung | 10X | *Rpl37a,Rps12,Rps15,Rpl41,Rps27a,Rpl23a,Rpl23,Rpl19,Rpl38,Rps7,Rps29,Rps23,Rps24,Tpt1,Rpl37,Rpl8,Rpl35a,Rps18,Rpl36,Rps14,Rpl17,Rpl35,Rps3a,Rps8,Rps15a-ps4,Rpl11,Rpl9,Rpl6,Rpl32,Rps5,Rps16,Rpl13a,Rps3,Rpl27a,Rps13,Rps15a,Rplp2,Uba52,Rpl18a,Rpl13,Rps25,Rplp1,Eef1a1,Rpl39,Rpl10,Rps4x* |
| Mammary_Gland | 10X | *Rpl37a,Rps15,Rps27a,Rpl23a,Rpl23,Rpl19,Rpl38,Rps7,Rps29,Rps23,Rps24,Rpl37,Rpl8,Rpl35a,Rps10,Rps18,Rpl36,Rps14,Rpl17,Fau,Rpl35,Rps3a,Rps20,Rps6,Rps8,Rpl11,Rpl9,Rplp0,Rpl6,Rpl21,Rpl32,Rps9,Rps5,Rps19,Rps16,Rpl13a,Rps3,Rpl27a,Rps13,Rps15a,Rplp2,Uba52,Rpl18a,Rpl13,Rps25,Rplp1,Eef1a1,Rpl14,Rpl39,Rps4x* |
| Marrow | 10X | *Rpl7,Rpl37a,Rps12,Rps15,Naca,Rpl41,Rps26,Rps27a,Gnb2l1,Rpl23a,Rpl23,Rpl19,Rpl27,Rpl38,Rps7,Rps29,Rps23,Rpl15,Rps24,Tpt1,Rpl37,Rpl8,Rpl3,Rpl35a,Rpl24,Rps2,Rps10,Rpl10a,Rps28,Rps18,Rpl36,Rps14,Rpl17,Fau,Rpl7a,Rpl35,Rps21,Rps3a,Rps20,Rps6,Rps8,Rpl11,Rpl22,Rpl9,Rpl5,Rplp0,Rpl6,Rpl21,Rpl32,Rps9,Rpl28,Rps5,Rps19,Rps16,Rps11,Rpl13a,Rpl18,Rps17,Rps3,Rpl27a,Rps13,Rps15a,Rplp2,Uba52,Rpl18a,Rpl13,Rps25,Rplp1,Rpl4,Eef1a1,Rpl29,Rpsa,Rpl14,Rpl39,Rpl10,Rps4x,Rpl36a* |
| Spleen | 10X | *Rpl37a,Rps15,Rps27a,Rpl23a,Rpl23,Rpl19,Rps7,Rps29,Rps23,Rps24,Rpl8,Rpl35a,Rps18,Rps14,Rpl17,Fau,Rpl35,Rps3a,Rps6,Rps8,Rpl11,Rpl9,Rpl6,Rpl21,Rpl32,Rps9,Rps5,Rps19,Rps16,Rpl13a,Rps3,Rpl27a,Rps13,Rps15a,Rplp2,Uba52,Rpl18a,Rpl13,Eef1a1,Rps4x* |
| Thymus | 10X | *Rpl7,Rpl31,Rpl37a,Rps15,Rpl41,Rps27a,Rpl23a,Rpl23,Rpl19,Rpl27,Rps7,Rps29,Rps23,Rpl15,Rps24,Tpt1,Rpl37,Rpl8,Rpl35a,Rpl24,Rps10,Rpl36,Rps14,Rpl17,Fau,Rpl35,Rps3a,Rps6,Rps8,Rps15a-ps4,Rpl11,Rpl9,Rplp0,Rpl6,Rpl21,Rpl32,Rps9,Rpl28,Rps5,Rps16,Rps11,Rpl13a,Rpl18,Rps17,Rps3,Rpl27a,Rps13,Rps15a,Rplp2,Uba52,Rpl18a,Rpl13,Rpl4,Eef1a1,Rpl39,Rpl10,Rps4x* |
| Tongue | 10X | *Rpl7,Rpl31,Eef1b2,Rpl37a,Rps12,Rps15,Eef2,Naca,Rpl41,Rps26,Rps27a,Gnb2l1,Rpl23a,Rpl23,Rpl19,Rpl27,Rpl38,Rps7,Rps29,Rpl36al,Rps23,Btf3,Rpl15,Rps24,Tpt1,Rpl37,Eif3h,Rpl8,Rpl3,Rpl35a,Rpl24,Rps2,Rps10,Rpl10a,Rps28,Rps18,Rpl36,Rps14,Rpl17,Fau,Eef1g,Rpl7a,Rpl12,Rpl35,Rps21,Rpl22l1,Rps3a,Rps27,Rps20,Rps6,Uqcrh,Rps8,Rpl11,Rpl22,Rpl9,Rpl5,Rplp0,Rpl6,Rpl21,Rpl32,Rps9,Rpl28,Rps5,Rps19,Rps16,Rps11,Rpl13a,Rpl18,Rps17,Rps3,Eif3f,Rpl27a,Rps13,Rps15a,Rplp2,Uba52,Rpl18a,Cox4i1,Rpl13,Rps25,Rplp1,Rpl4,Eef1a1,Rpl29,Rpsa,Rpl14,Rpl39,Rpl10,Rps4x,Rpl36a* |
| Trachea | 10X | *Rpl7,Rpl37a,Rps12,Rps15,Rpl41,Rps27a,Rpl23a,Rpl23,Rpl19,Rps7,Rps29,Rps23,Rps24,Tpt1,Rpl37,Rpl8,Rpl35a,Rps2,Rps18,Rps14,Rpl17,Rpl35,Rps3a,Rps6,Rps8,Rpl11,Rpl9,Rplp0,Rpl6,Rpl21,Rpl32,Rps9,Rps5,Rps19,Rps16,Rpl13a,Rps3,Rpl27a,Rplp2,Rpl18a,Rpl13,Rplp1,Eef1a1,Rpl14,Rpl10,Rps4x* |

#### 3.4 Supplementary table S4

**Table S4 The computational efficiency of ISnorm on two datasets from 10X Genomics Chromium system.**

| dataset | number of cells | number of cells passing quality control | number of input genes (detection rate) | running time |
| --- | --- | --- | --- | --- |
| mammary epithelial cells (16) | 25,806 | 25,114 | 172 (90%) | 2.80 mins |
|  |  |  | 480 (70%) | 11.49 mins |
| PBMCs | 11,769 | 10,759 | 179 (90%) | 1.77 mins |
|  |  |  | 407 (70%) | 5.12 mins |

### **4. Supplementary Note**

#### **4.1 Supplementary Note 1**

Pearson correlation coefficient (PCC) is widely used in biology to measure the linear relationship between genes. However, previous study has considered that it is not appropriate for compositional data (Chapter3 in (20)). Specially, it does not maintain sub compositional coherence for scRNA-seq data. We showed *DoR* provided more robust estimate of linear relationship than PCC by one scRNA-seq data of mouse embryonic stem cells (ESCs) in figure below (12). This dataset contains UMI (unique molecule identifier) counts of 933 mouse ESCs generated by inDrop protocol. First we calculated PCC (left below) and *DoR* (right below) between a set of constantly expressed genes on the full dataset of 933 cells (these genes are selected by ISnorm and should have good linear relationship with each other). As expected, PCC scores between IS genes were high (all PCC scores are above 0.5, red dots). Then we calculated PCC and *DoR* between these genes on a partial dataset by subsampling cells with library size larger than 20,000 but smaller than 30,000. We found that most PCC scores decreased significantly (mostly below 0.5 on the partial dataset, compared red and blue dots in left panel) but *DoR* roughly remained the same (compared red and blue dots in right panel). We also calculated PCC between these genes on the full dataset but scaling the library size of each cell to be the same to mimic the case of equal sequencing depth for all cells. Similar decrease in PCC can be observed (green dots in left panel) while *DoR* is not affected by such scaling.

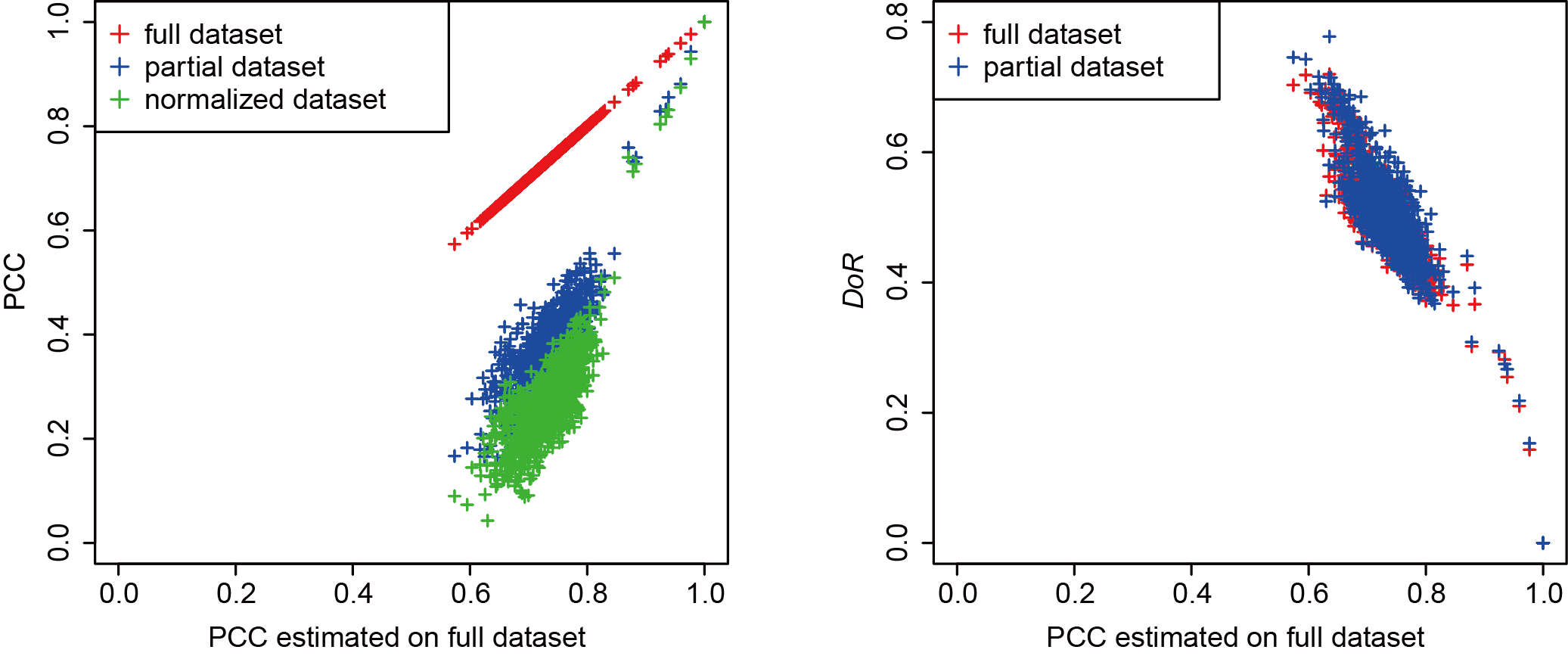

The *DoR* statistic is a modified version of log-ratio variance (LRV) of two gene vectors (20), with the mean value replaced by the median value. Assuming that gene ***X*** and gene ***Y*** are constantly expressed in most cells but differentially expressed in some rare cell types, most log ratios (*Z*) will be close except for few outliers. In this case, *DoR* centered on the median value will provide a larger distance measurement than the *SD* of ***Z***, which can help prevent the information of rare cell types from being overwhelmed by the majority of cell population.

In Erb & Notredame paper (21) the authors also proposed a modified version of LRV scaled by the sum of log variance of two genes. Such scaling may help find co-regulated patterns between two genes. Based on our test, *DoR* and scaled LRV performed similarly in most cases. But *DoR* worked better when there were drastic changes in global transcriptome, and we suspected that the scaling by sum of log variance might preferentially select the lowly expressed genes as stably expressed genes.

Finally, the *DoR* between two genes or two cells remains unchanged when we scale the expression of each cell or each gene by a constant (e.g. size factor for cells and/or effective length for genes). This feature enables ISnorm to re-normalize most normalized matrices, such as a FPKM or TPM matrix, which is easier to obtain from public database.

#### **4.2 Supplementary Note 2**

As described by (9), single cells were sorted into 96-well plate. If we assume each well had similar reverse-transcription and PCR efficiency, more cDNAs would be obtained from cells with more starting mRNA materials. By pooling 96 cells together before sequencing, more reads would be generated from cells with more cDNAs. Thus we hypothesize that library size may reflect the total mRNA content of each cell. The high consistency between normalized mRNA content upon ISnorm and library size supports our hypothesis (Supplementary Fig. 6C).

#### **4.3 Supplementary Note 3**

We noticed that the threshold based approach to select IS genes performed well in deep sequencing data but reported few IS genes in some shallow sequencing data, indicating that sequencing uncertainty could also cause poor linear relationships between genes. In order to account for sequencing depth, we developed a method to generate an empirical null distribution for the instability scores and estimate the *p* values of candidate genesets. Given one candidate geneset containing *n* IS genes with reference expression $\boldsymbol{REF}={({ref}_{1}, {ref}_{2}, \ldots\ldots, {ref}_{n})}^{T}$ and cell *j* with size factor *sf_j_*, the expected expression of IS genes in cell *j* were defined as follows:

***Ê_j_*** = *sf_j_ ∙* ***REF***

where ***Ê_j_*** = (*ê_1j_*, *ê_2j_*, ... *ê_nj_*)*^T^*. Then we sampled the raw counts of gene *i* in cell *j* from a Poisson distribution with parameter *λ_ij_* = *ê_ij_* for 1,000 times and generated 1,000 simulated cells for cell *j*. We calculated the instability scores of simulated cells and generated an empirical null distribution. Finally we compared the observed instability score of cell *j* with the null distribution. This process assigned a right-tail *p* value for each cell given the candidate geneset. If the percentage of cells with *p* < 0.05 was far higher than 5%, the candidate geneset was less likely to be true IS genes. For deep sequencing data, as we did not introduce biological variations, this strategy would generate extremely low instability scores, which did not agree with real datasets. But the threshold based approach could control the biological variation of IS genes well in these cases.

We doubted that the high instability scores of non-optimal genesets in Zheng dataset might derive from sequencing uncertainty and tested our hypothesis through the strategy described above. We found that the non-optimal genesets in P0205 and P0508 cannot be explained by sequencing uncertainty, while the optimal geneset did not show strong deviance from the null distribution (see the table below and Supplementary Figure 9). As the non-optimal genesets in P0205 and P0508 could not pass either threshold based or null distribution based tests, they were less likely to be true IS genes.

| Patient | IS genes | Median instability scores | Mean instability scores | Percentage of cells with *p* < 0.05 |
| --- | --- | --- | --- | --- |
| P0205* | 5 IS genes: *HLA-A, HLA-B, HLA-G, HLA-H, HLA-J* | 0.095 | 0.111 | 1.2% |
|  | 12 IS genes: *RPL30, RPL27A, RPL31, RPL32, RPL34, RPL37, RPL37A, RPLP1, RPLP2, RPS14, RPS18, RPS29* | 0.159 | 0.175 | 68.2% |
|  | 16 IS genes: *RPL11, RPL19, RPL30, RPL27A, RPL31, RPL32, RPL34, RPL37, RPL37A, RPLP1, RPLP2, RPS14, RPS18, RPS27A, RPS29, RPS14P3* | 0.171 | 0.187 | 40.9% |
| P0322 | 5 IS genes: *HLA-A, HLA-B, HLA-G, HLA-H, HLA-J* | 0.079 | 0.089 | 2.6% |
|  | 11 IS genes: *CFL1, HLA-A, HLA-B, HLA-C, HLA-E, HLA-G, HLA-H, HLA-J, HLA-L, B2M, ACTB* | 0.163 | 0.174 | 6.2% |
|  | 15 IS genes: *CFL1, HLA-A, HLA-B, HLA-C, HLA-E, HLA-G, HLA-H, HLA-J, HLA-L, MALAT1, PFN1, B2M, SPPL2B, PTPRC, ACTB* | 0.200 | 0.213 | 16.1% |
| P0407 | 5 IS genes: *HLA-B, HLA-C, HLA-G, HLA-L, B2M* | 0.123 | 0.143 | 1.4% |
|  | 10 IS genes: *CFL1, HLA-A, HLA-B, HLA-C, HLA-E, HLA-G, HLA-H, HLA-L, MALAT1, B2M* | 0.176 | 0.193 | 5.5% |
|  | 17 IS genes: *CFL1, HLA-A, HLA-B, HLA-C, HLA-E, HLA-G, HLA-H, HLA-J, HLA-L, MALAT1, B2M, PTPRC, ACTB, RPL13, RPS3, RPS6, TMSB4X* | 0.218 | 0.238 | 22.7% |
| P0508* | 5 IS genes: *HLA-A, HLA-B, HLA-G, HLA-H, HLA-J* | 0.074 | 0.084 | 1.9% |
|  | 10 IS genes: *RPL13, RPL27A, RPL34, RPL37, RPLP2, RPS3, RPS14, RPS27A, RPS29, RPS14P3* | 0.167 | 0.182 | 69.1% |
|  | 15 IS genes: *RPL13, RPL30, RPL27A, RPL31, RPL34, RPL37, RPLP1, RPLP2, RPS3, RPS14, RPS18, RPS25, RPS27A, RPS29, RPS14P3* | 0.182 | 0.193 | 87.7% |
| P1116 | 5 IS genes: *HLA-A, HLA-B, HLA-G, HLA-H, HLA-J* | 0.083 | 0.098 | 15.0% |
|  | 10 IS genes: *HLA-A, HLA-B, HLA-C, HLA-G, HLA-H, HLA-J, HCG4, LOC554223, B2M, ACTB* | 0.145 | 0.167 | 11.6% |
|  | 16 IS genes: *CFL1, HLA-A, HLA-B, HLA-C, HLA-E, HLA-G, HLA-H, HLA-J, MALAT1, ARHGDIB, HCG4, LOC554223, B2M, ACTB, TMSB4X, IL32* | 0.213 | 0.235 | 58.1% |

### 5. References

1.	Deng, Q., Ramskold, D., Reinius, B. and Sandberg, R. (2014) Single-cell RNA-seq reveals dynamic, random monoallelic gene expression in mammalian cells. *Science (New York, N.Y.)*, **343**, 193-196.

2.	Yan, L., Yang, M., Guo, H., Yang, L., Wu, J., Li, R., Liu, P., Lian, Y., Zheng, X., Yan, J. *et al.* (2013) Single-cell RNA-Seq profiling of human preimplantation embryos and embryonic stem cells. *Nature structural & molecular biology*, **20**, 1131-1139.

3.	Goolam, M., Scialdone, A., Graham, S.J.L., Macaulay, I.C., Jedrusik, A., Hupalowska, A., Voet, T., Marioni, J.C. and Zernicka-Goetz, M. (2016) Heterogeneity in Oct4 and Sox2 Targets Biases Cell Fate in 4-Cell Mouse Embryos. *Cell*, **165**, 61-74.

4.	Love, M.I., Huber, W. and Anders, S. (2014) Moderated estimation of fold change and dispersion for RNA-seq data with DESeq2. *Genome Biol*, **15**, 550.

5.	Wu, J., Huang, B., Chen, H., Yin, Q., Liu, Y., Xiang, Y., Zhang, B., Liu, B., Wang, Q., Xia, W. *et al.* (2016) The landscape of accessible chromatin in mammalian preimplantation embryos. *Nature*, **534**, 652-657.

6.	Li, H., Handsaker, B., Wysoker, A., Fennell, T., Ruan, J., Homer, N., Marth, G., Abecasis, G. and Durbin, R. (2009) The Sequence Alignment/Map format and SAMtools. *Bioinformatics (Oxford, England)*, **25**, 2078-2079.

7.	Robinson, J.T., Thorvaldsdottir, H., Winckler, W., Guttman, M., Lander, E.S., Getz, G. and Mesirov, J.P. (2011) Integrative genomics viewer. *Nature biotechnology*, **29**, 24-26.

8.	Ramirez, F., Dundar, F., Diehl, S., Gruning, B.A. and Manke, T. (2014) deepTools: a flexible platform for exploring deep-sequencing data. *Nucleic acids research*, **42**, W187-191.

9.	Patel, A.P., Tirosh, I., Trombetta, J.J., Shalek, A.K., Gillespie, S.M., Wakimoto, H., Cahill, D.P., Nahed, B.V., Curry, W.T., Martuza, R.L. *et al.* (2014) Single-cell RNA-seq highlights intratumoral heterogeneity in primary glioblastoma. *Science (New York, N.Y.)*, **344**, 1396-1401.

10.	Subramanian, A., Tamayo, P., Mootha, V.K., Mukherjee, S., Ebert, B.L., Gillette, M.A., Paulovich, A., Pomeroy, S.L., Golub, T.R., Lander, E.S. *et al.* (2005) Gene set enrichment analysis: a knowledge-based approach for interpreting genome-wide expression profiles. *Proceedings of the National Academy of Sciences of the United States of America*, **102**, 15545-15550.

11.	Tang, F., Barbacioru, C., Wang, Y., Nordman, E., Lee, C., Xu, N., Wang, X., Bodeau, J., Tuch, B.B., Siddiqui, A. *et al.* (2009) mRNA-Seq whole-transcriptome analysis of a single cell. *Nature methods*, **6**, 377-382.

12.	Klein, A.M., Mazutis, L., Akartuna, I., Tallapragada, N., Veres, A., Li, V., Peshkin, L., Weitz, D.A. and Kirschner, M.W. (2015) Droplet barcoding for single-cell transcriptomics applied to embryonic stem cells. *Cell*, **161**, 1187-1201.

13.	Leng, N., Chu, L.F., Barry, C., Li, Y., Choi, J., Li, X., Jiang, P., Stewart, R.M., Thomson, J.A. and Kendziorski, C. (2015) Oscope identifies oscillatory genes in unsynchronized single-cell RNA-seq experiments. *Nature methods*, **12**, 947-950.

14.	Nestorowa, S., Hamey, F.K., Pijuan Sala, B., Diamanti, E., Shepherd, M., Laurenti, E., Wilson, N.K., Kent, D.G. and Gottgens, B. (2016) A single-cell resolution map of mouse hematopoietic stem and progenitor cell differentiation. *Blood*, **128**, e20-31.

15.	Zheng, C., Zheng, L., Yoo, J.K., Guo, H., Zhang, Y., Guo, X., Kang, B., Hu, R., Huang, J.Y., Zhang, Q. *et al.* (2017) Landscape of Infiltrating T Cells in Liver Cancer Revealed by Single-Cell Sequencing. *Cell*, **169**, 1342-1356.e1316.

16.	Bach, K., Pensa, S., Grzelak, M., Hadfield, J., Adams, D.J., Marioni, J.C. and Khaled, W.T. (2017) Differentiation dynamics of mammary epithelial cells revealed by single-cell RNA sequencing. *Nat Commun*, **8**, 2128.

17.	(2018) Single-cell transcriptomics of 20 mouse organs creates a Tabula Muris. *Nature*, **562**, 367-372.

18.	Ziegenhain, C., Vieth, B., Parekh, S., Reinius, B., Guillaumet-Adkins, A., Smets, M., Leonhardt, H., Heyn, H., Hellmann, I. and Enard, W. (2017) Comparative Analysis of Single-Cell RNA Sequencing Methods. *Molecular cell*, **65**, 631-643.e634.

19.	Schep, A.N., Wu, B., Buenrostro, J.D. and Greenleaf, W.J. (2017) chromVAR: inferring transcription-factor-associated accessibility from single-cell epigenomic data. *Nature methods*, **14**, 975-978.

20.	Aitchison, J. (1986) *The Statistical Analysis of Compositional Data*.

21.	Erb, I. and Notredame, C. (2016) How should we measure proportionality on relative gene expression data? *Theory in biosciences = Theorie in den Biowissenschaften*, **135**, 21-36.
